# Microglia Clear Neuron-released α-Synuclein via Selective Autophagy and Prevent Neurodegeneration

**DOI:** 10.1101/2019.12.11.872812

**Authors:** Insup Choi, Yuanxi Zhang, Steven P. Seegobin, Mathilde Pruvost, Qian Wang, Kerry Purtell, Bin Zhang, Zhenyu Yue

## Abstract

Microglia maintain brain homeostasis by removing neuron-derived components such as myelin and cell debris. The evidence linking microglia to neurodegenerative diseases is growing, however, the precise mechanisms remain poorly understood. Herein we report a neuroprotective role for microglia in the clearance of neuron-released α-synuclein. Neuronal α-synuclein activates microglia, which in turn engulf α-synuclein into autophagosomes for degradation via selective autophagy (termed Synucleinphagy). Synucleinphagy is initiated by the interaction of α-synuclein with Toll-like receptor 4 (TLR4), which induces transcriptional upregulation of *p62/SQSTM1* through the NF-κB signaling pathway without causing TLR4 endocytosis. Induction of p62, an autophagy receptor, is necessary for the formation of α-synuclein/ubiquitin-positive puncta that are degraded by autophagy. Finally, disruption of microglial autophagy in mice expressing human α-synuclein promotes the accumulation of misfolded α-synuclein and causes midbrain dopaminergic neuron degeneration. Our study thus identifies a novel neuroprotective function of microglia in the clearance of α-synuclein via TLR4-NF-κB-p62 mediated Synucleinphagy.

## Introduction

α-synuclein is produced primarily in neurons and constitutes up to 1% of total cytosolic protein in the brain^1^. Although the function of α-synuclein is poorly understood, available evidence suggests its role in synaptic vesicular trafficking^2–4^. Aggregated α-synuclein is a major component in intraneuronal inclusions known as Lewy bodies (LB) associated with neurodegenerative diseases such as Parkinson’s disease (PD) and dementia with Lewy body (DLB)^5, 6^. Increased α-synuclein level due to the multiplication of *SNCA* alleles is causal to PD^7, 8^.

Previous evidence suggested “prion-like” cell-to-cell transmission of α-synuclein^9, 10^. α-synuclein can be secreted by neurons as a result of cellular stress or injury, or as a response to stimulation^11–13^. Neighboring neurons and glia can engulf and clear extracellular α-synuclein and thus contributing to the regulation of α-synuclein homeostasis in the brain. Interestingly, injection of fibrillar α-synuclein into animal brains causes the spread of LB-like pathology^14^, supporting the “Braak hypothesis” of staging in synucleinopathies^15^. Therefore, cellular uptake and clearance pathways are key processes to control the deposition and spread of α-synuclein aggregates, thus affecting disease progression.

Although neurons and glia in the brain can ingest and degrade extracellular α-synuclein, microglia show the highest efficiency *in vitro*^16^. Extensive effort has been made towards the identification of cellular pathways of α-synuclein clearance. Several studies reported receptor-mediated internalization of various forms of α-synuclein in neurons and glial cells^17–19^. However, the exact mechanism for the clearance of internalized α-synuclein remains unclear.

Previous studies suggest that α-synuclein is degraded by macroautophagy, chaperone-mediated autophagy, and the proteasome^20–23^. However, the evidence for macroautophagy (hereafter referred to as autophagy) degradation of α-synuclein is extremely limited^24, 25^. Autophagy is a bulk degradation pathway responsible for the clearance of protein aggregates and damaged cellular organelles^26, 27^. Recent evidence demonstrates a strong selectivity for autophagy^28^. Direct evidence for autophagy in selective degradation of α-synuclein is entirely lacking. Furthermore, since most of the studies of α-synuclein degradation were focused on neurons, whether or not microglia, the prototypical scavenger cell in the brain, take part in the degradation of α-synuclein remains unclear.

Here we report that microglia ingest and degrade neuron-released α-synuclein through selective autophagy *in vitro* and *in vivo*. We document that ingested α-synuclein in microglia is sequestered by autophagosomes for degradation, which is mediated by TLR4-NF-κB signaling through upregulation of the autophagy receptor, *p62/SQSTM1*. Thus, our results uncover a microglia-specific α-synuclein degradation pathway termed *Synucleinphagy* that regulates α-synuclein homeostasis in the CNS.

## Results

### Neuron-released α-synuclein induces microglial activation and is engulfed by microglia *in vivo*

To understand microglia and α-synuclein interaction *in vivo*, we employed two mouse models, both of which express wild-type (WT) human α-synuclein (*h*α-Syn) in a neuron-specific manner. We first verified that mice injected with adeno-associated virus 9 (AAV9) carrying green fluorescent protein (GFP) under the synapsin promoter into the Substantia Nigra pars compacta (SNpc) expressed GFP strictly in neurons (Supplementary Fig. 1a, b). Mice injected with AAV9-*h*α-Syn in the SNpc showed loss of dopaminergic neurons on the ipsilateral side (Supplementary Fig. 1c) as reported^29^. In fact, dopamine transporter (DAT) and tyrosine hydroxylase (TH) protein levels were reduced compared to AAV9-GFP injected mice (Supplementary Fig. 1d, e). Moreover, AAV9-*h*α-Syn, but not AAV9-GFP injection, caused microglial activation in the striatum as evidenced by the reduced length of processes and terminal branch points as well as increased cell number (Fig. 1a, b).

**Figure 1.**
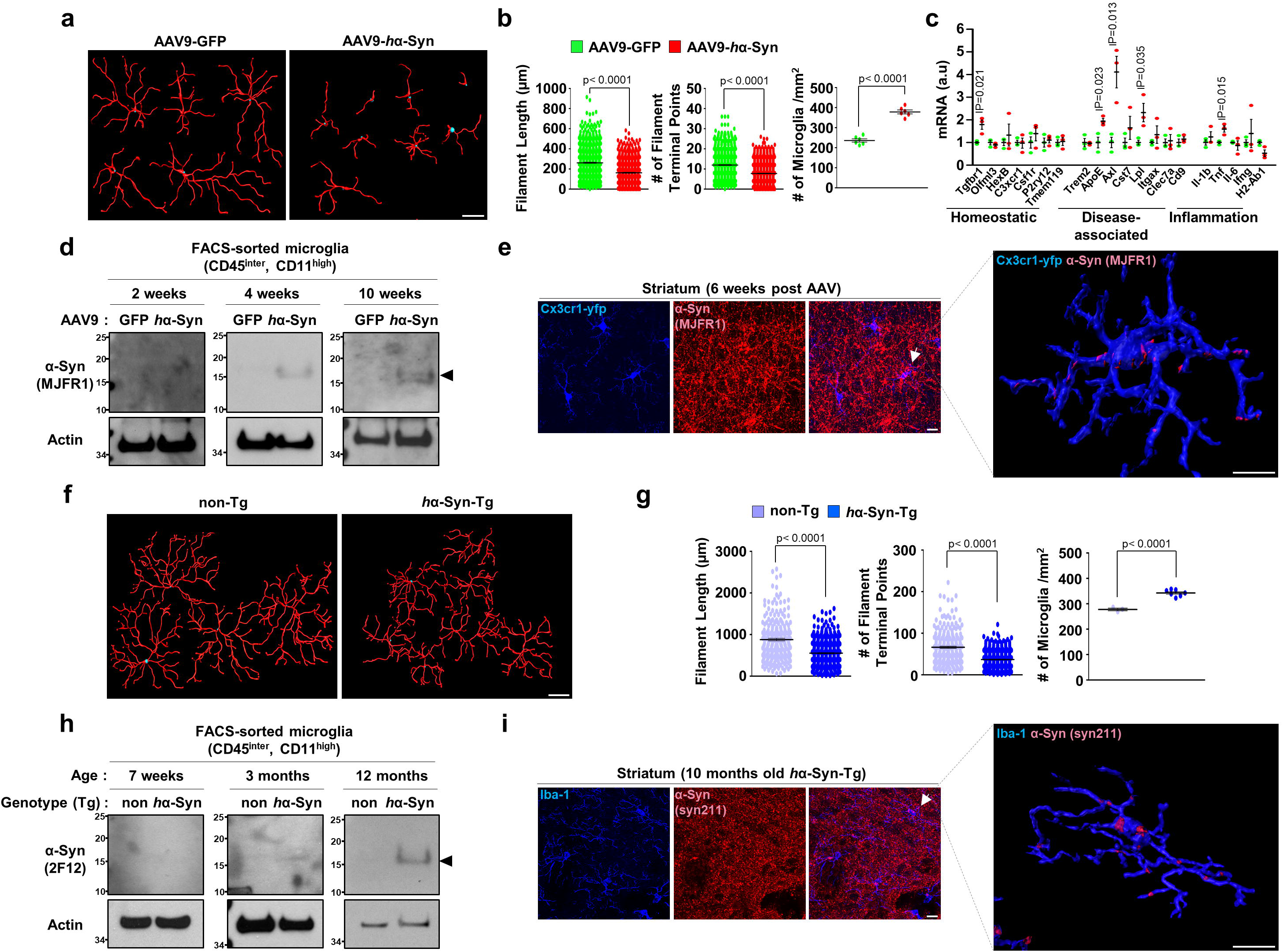
Neuron-released α-synuclein induces microglial activation and is engulfed by microglia *in vivo*. **(a, b, f, g)** Brain sections from AAV-GFP (n=6) and AAV-*h*α-Syn-injected mice (n=6) at 6-weeks post AAV injection (**a, b**) or from 10-months-old Non-Tg (n=3) and *h*α-Syn-Tg (n=7) mice (**f, g**) were stained for a marker of microglia, Iba-1, to visualize microglia morphology in the striatum. Reconstruction of microglia processes was generated by Filament-Tracer tool in Imaris software (a, e, Bitplane) and filament lengths and the number of terminal points were quantified (left panels in b and g). The number of microglia at the striatum were counted using ImageJ software (NIH, Bethesda, MD) (right panel in b and g). *p* values were calculated by Mann-Whitney U test (the left two panels) and Unpaired two-tailed Student’s t-test (the right panel). Scale bar, 10 µm. **(c)** CD45^intermediate^ and CD11b^high^ microglia were isolated from brains injected with AAV-*h*α-Syn (n=3) and assayed for RT-qPCR using primers as indicated, compared to AAV-GFP (n=3) at 4-weeks after AAV. *p* values were calculated by Unpaired two-tailed Student’s t-test. **(d, h)** Pooled CD45^intermediate^ and CD11b^high^ microglia were prepared either from AAV-GFP (n=3) and AAV-*h*α-Syn-injected mice (n=3) (**d**) or Non-Tg (n=4) and *h*α-Syn-Tg (n=4) mice (**h**) brains at the indicated time points and lysed for W.B using α-synuclein antibodies (MJFR1 clone in d, 2F12 clone in h) and actin. **(e, i)** Brain sections from *Cx3cr1*^CreER-IRES-Eyfp^ mice injected with AAV-*h*α-Syn were stained using anti-GFP/YFP antibody and human α-synuclein antibody at 6-weeks after AAV (**e**), and brain sections from 10-months-old Non-Tg and α-Syn-Tg mice were stained using Iba-1 antibody and human α-Synuclein antibody (**i**). 3D reconstruction of microglia and human α-synuclein was produced using Imaris software (Bitplane, right panel) as described in Method. Scale bar, 10µm. All values are reported as mean ± SEM.

By RT-qPCR analysis of microglia state-specific gene expression^30–32^, we found upregulation of several disease-associated signature genes such as *ApoE*, *Axl*, and *Lpl*, inflammatory gene *Tnf*, and *Tgfbr1* at 4-weeks post-AAV injection (Fig. 1c and Supplementary Fig. 2). Importantly, we verified the presence of *h*α-Syn protein in isolated microglia via Western blot with *h*α-Syn specific antibody (Fig. 1d and Supplementary Fig. 3a). Using *Cx3cr1*^CreER-IRES-Eyfp^ transgenic mice^33^, which express enhanced yellow fluorescent protein (EYFP) under the microglia-specific *Cx3cr1* promoter, we detected *h*α-Syn in Cx3cr1-EYFP-positive microglia (Fig. 1e and Supplementary Fig. 3b).

In transgenic mice overexpressing *h*α-Syn under the Thy-1 promoter (*h*α-Syn-Tg), we also found microglial activation in the striatum of 10-months-old mice (Fig. 1f, g). *h*α-Syn protein was clearly present in microglia through Western blot analysis (Fig. 1h) and immunofluorescent staining in Iba1-positive microglia at the striatum (Fig. 1i and Supplementary Fig. 3c). Noticeably, *h*α-Syn-Tg mice did not show degeneration of dopaminergic neurons in SNpc up to 12-months-old (Supplementary Fig. 1f). Together, our data from two different mouse models demonstrate that neuron-released *h*α-Syn induces microglial activation, accompanied by microglial engulfment of extracellular *h*α-Syn *in vivo*.

### Microglia-engulfed α-synuclein is sequestered by autophagosomes and degraded by autophagy

We next asked if ingested α-synuclein goes through endocytic degradation following phagocytosis or receptor-mediated endocytosis^17, 19^. We treated cultured primary microglia with recombinant *h*α-Syn protein and found no evidence of colocalization between *h*α-Syn and an early endosome marker EEA1 (Supplementary Fig. 3d and 4a). Neither cytochalasin D, a blocker of actin polymerization necessary for phagocytosis, nor Dynasore, a GTPase inhibitor that suppresses both clathrin-dependent and independent endocytosis, inhibited the uptake of *h*α-Syn protein but rather it increased the uptake (Supplementary Fig. 4b, 5a, 5b).

We then asked whether autophagy degrades *h*α-Syn in microglia. We co-stained *h*α-Syn-treated primary microglia with anti*-h*α-Syn and anti-ubiquitin or anti-p62/SQSTM1, an autophagy receptor^34^, and observed colocalization of *h*α-Syn with ubiquitin or p62 in discrete puncta (middle panels, Fig. 2a; left panels, Fig. 2b). We also treated microglia derived from GFP-LC3 transgenic mice^35^ with *h*α-Syn and detected co-localization of the autophagosome marker, GFP-LC3, and *h*α-Syn (right panels, Fig. 2b). These *h*α-Syn-associated ubiquitin-, GFP-LC3-, or p62-positive puncta appeared at 6 hours and disappeared at 24 hours after *h*α-Syn treatment (arrowheads, Fig. 2a). We also noticed smaller ubiquitin-negative *h*α-Syn particles that were evident after 6 hours but remained 24 hours after treatment (arrows, Fig. 2a). A time-course analysis showed that *h*α-Syn/ubiquitin-positive puncta emerged at 3 hours, and the number of puncta peaked at 6 hours, followed by a reduction in number at 24 hours post-treatment (left panel, Fig. 2c). The puncta number also depended on *h*α-Syn concentration (right panel, Fig. 2c). Additionally, we also observed *h*α-Syn-associated ubiquitin-, GFP-LC3-positive puncta after treating the medium from AAV9-*h*α-Syn-infected cortical neuron (arrowheads, Supplementary Fig. 4c), suggesting that *h*α-Syn-associated LC3 puncta also occurs in microglia in response to physiological form of α-Syn. Furthermore, in GFP-LC3 transgenic mice injected with AAV9-*h*α-Syn, we found ingested *h*α-Syn co-localized with GFP-LC3 puncta in Iba1-positive microglia in the striatum (Fig. 2d). We then performed ultrastructural analysis through electron microscopy (EM) and found that *h*α-Syn-treated microglia develop large inclusions which are surrounded by double-membrane structures, characteristic of autophagosomes (Fig. 2e).

**Figure 2.**
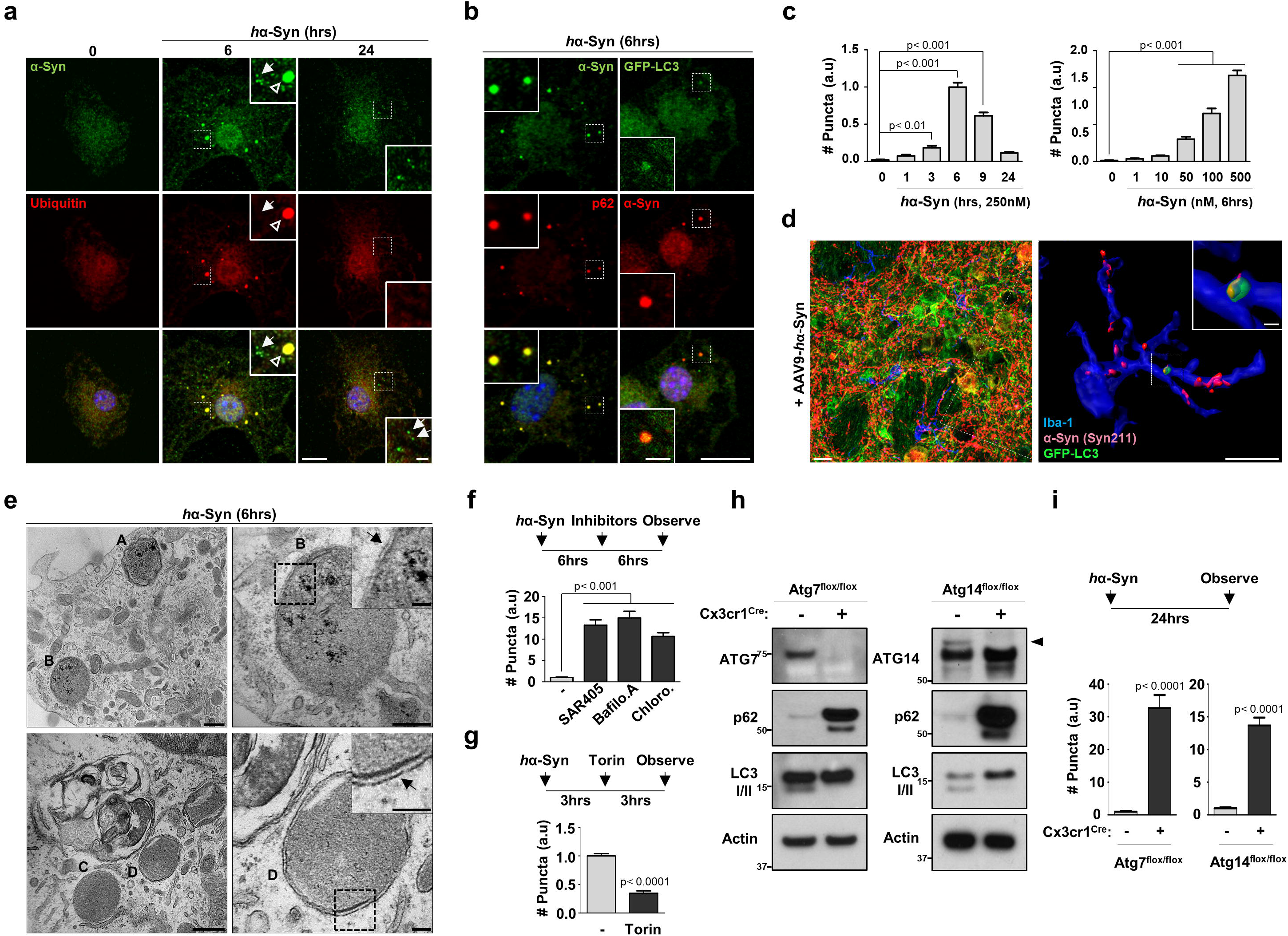
Microglia-engulfed α-synuclein is sequestered by autophagosomes and degraded by autophagy. **(a, b, c)** Cultured primary microglia from WT mice (a and left panels of b) and GFP-LC3-transgenic mice (right panels of b) were treated 250nM *h*α-Syn protein for indicated time and stained with antibodies against human α-synuclein (MJFR1 clone, a and b), ubiquitin (**a**), or p62 (**b**). The number of *h*α-Syn/ubiquitin-positive puncta was quantified (**c**). In (a), empty arrowheads indicate a portion of α-synuclein colocalized with ubiquitin-positive puncta and disappear in 24 hours. Arrows represent a portion of α-synuclein not colocalized with ubiquitin. *p* values were calculated by One-way ANOVA with Newman–Keuls post hoc test. Scale bar, 10 µm; 2 µm in magnified boxes. **(d)** At 6-weeks post-AAV-*h*α-Syn inoculation into GFP-LC3 mice, brain slices were fixed and stained with antibodies against human α-synuclein, GFP, and Iba-1 (left panel). 3D surface-rendering images of each protein in microglia were made with Imaris software (Bitplane, right panel). Scale bar, 10µm; 1µm in the magnified box. **(e)** Ultrastructure of puncta from 250nM *h*α-Syn protein-treated cells for 6 hours was observed by an electron microscope. Arrows indicate double-membrane structures. Scale bar, 500 nm in the left panel; 100 nm in the right panel. **(f, g)** Microglia were treated with 250nM *h*α-Syn protein for 6 hours (f) or 3 hours (g) first and added SAR405 (**f**) or Torin (**g**) was added. The number of *h*α-Syn/ubiquitin-positive puncta was quantified in the lower panel. *p* values were calculated by One-way ANOVA with Newman– Keuls post hoc test (f) and Unpaired two-tailed Student’s t-test (g). **(h, i)** Microglia cultured either from *Atg7*^flox/flox^ mice (left panel) from *Atg14*^flox/flox^ mice (right panel) with or without *Cx3cr1^Cre^* expression were assayed for W.B using antibodies against ATG7 or ATG14, p62, and LC3 I/II (**h**). Cells were treated with 250nM *h*α-Syn protein for 24 hours and the number of *h*α-Syn/ubiquitin-positive puncta was quantified (**i**). *p* values were calculated by Mann-Whitney U test. Data are representative of at least three independent experiments.

We next treated microglia with SAR405, an inhibitor of VPS34, which is a class III phosphatidylinositol 3-kinase required for autophagosome membrane synthesis^36^, Bafilomycin A1, or Chloroquine, two different inhibitors of lysosome acidification, after adding *h*α-Syn. All three drugs prevented the clearance of *h*α-Syn/ubiquitin-positive puncta (Fig. 2f). In contrast, treatment with Torin1, an inhibitor of mTOR that stimulates autophagy^37^, facilitated the clearance of puncta (Fig. 2g). We next obtained autophagy-deficient microglia by breeding *Cx3cr1*^Cre^ mice, which constitutively express Cre recombinase, with either *Atg7*^flox/flox^ or *Atg14*^flox/flox^ mice. *Atg7* encodes an E1-like enzyme essential in the ubiquitin-like conjugation systems required for autophagy, and *Atg14* encodes a positive regulator of VPS34^38, 39^. The lack of autophagy was confirmed in *Atg7*-deficient cells or *Atg14*-deficient cells through the analysis of Atg7, Atg14, p62, and LC3-II (Fig. 2h). In these autophagy-deficient microglia, degradation of *h*α-Syn/ubiquitin-positive puncta was inhibited (Fig. 2i). We noticed that *Atg7* or *Atg14* deficiency did not cause microglia death regardless of *h*α-Syn protein treatment (Supplementary Fig. 4d, e). Thus, we concluded that microglia engulf extracellular α-syn protein, which is sequestered by autophagosomes and degraded by autophagy-lysosome pathway.

### Transcriptional upregulation of autophagy receptor p62 mediates microglial sequestration and degradation of α-synuclein

p62 is known to recognize and interact with ubiquitinated cargo to initiate selective autophagy^34^. We next investigated the role of p62 in autophagic clearance of *h*α-Syn. We observed that *h*α-Syn treatment caused an increase in p62 protein up to 9 hours (Fig. 3a). Interestingly, *p62* mRNA levels peaked at 3 hours, declined at 6 hours and returned to baseline at 9 hours post-treatment (Fig. 3b), which mirrors the transient increase of *h*α-Syn/ubiquitin/p62/GFP-LC3-positive puncta (Fig. 2a, b, c). Actinomycin D, a blocker of mRNA translation, inhibited p62 induction when added before *h*α-Syn treatment (Supplementary Fig. 4g) and suppressed the formation of *h*α-Syn/ubiquitin-positive puncta (Supplementary fig. 4h). In contrast, total LC3-II levels changed little after *h*α-Syn treatment (Supplementary Fig. 4i), suggesting no significant alteration of overall autophagy. Furthermore, other autophagy receptors such as Ndp52, Optineurin, and Nbr1 were not altered (Supplementary Fig. 4i), suggesting that *h*α-Syn selectively induces p62 expression without affecting other autophagy receptors.

**Figure 3.**
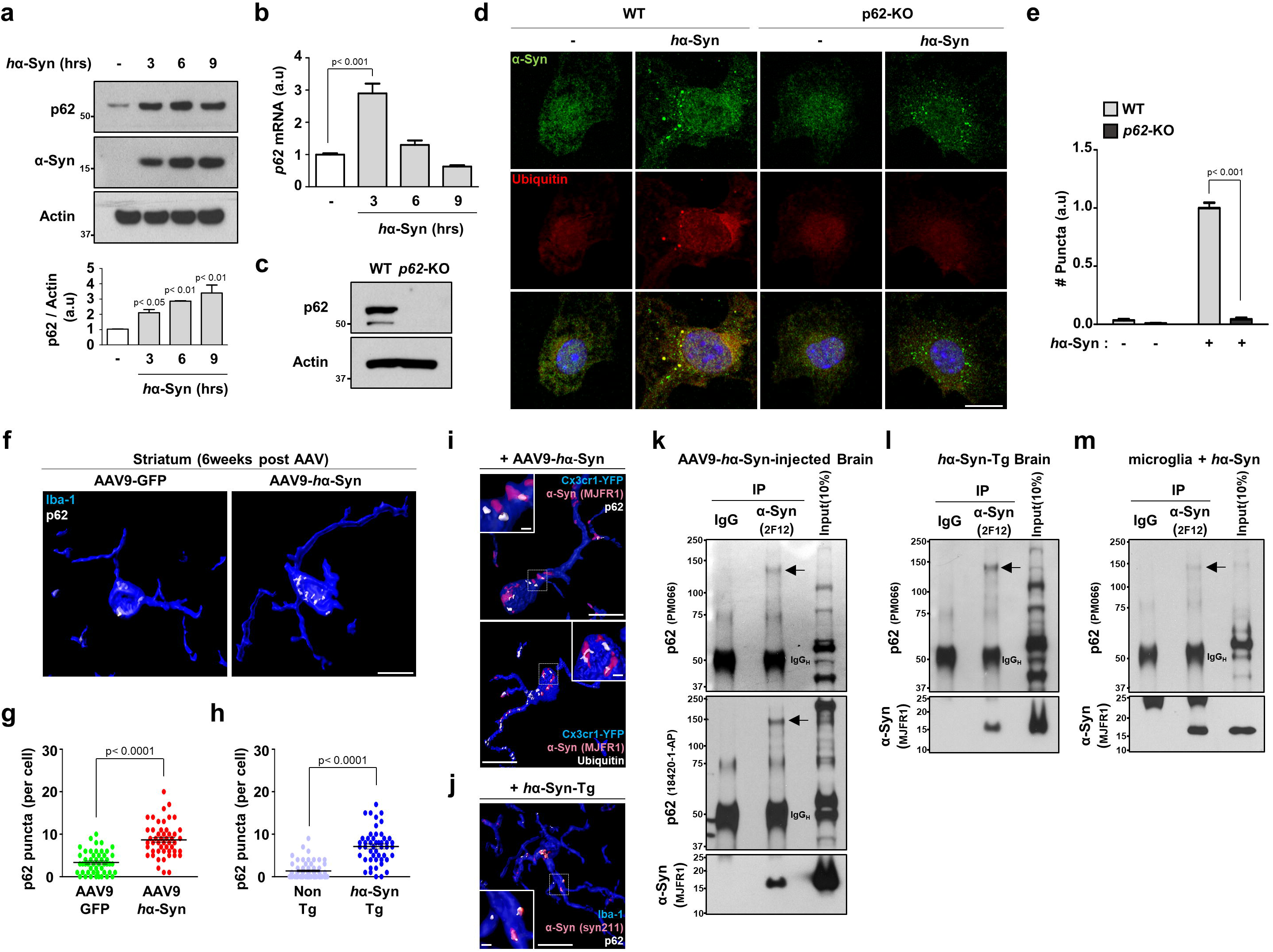
Transcriptional upregulation of autophagy receptor p62 mediates microglial sequestration and degradation of α-synuclein. **(a)** Microglia were treated with 250nM *h*α-Syn protein for the indicated time and assayed for W.B using antibodies against p62 and human α-synuclein. The levels of p62 protein were quantified in the lower panel. *p* values were calculated by One-way ANOVA with Newman– Keuls post hoc test. **(b)** After treatment of 250nM *h*α-Syn protein for the indicated time, the levels of *p62* mRNA were examined by RT-qPCR. *p62* mRNA was normalized to *Actin* mRNA. *p* values were calculated by One-way ANOVA with Newman–Keuls post hoc test. **(c)** The absence of p62 protein was confirmed in microglia obtained from *p62*-KO mice compared to WT mice by W.B. **(d, e)** WT and *p62*-KO microglia were treated with 250nM *h*α-Syn protein for 6 hours and stained with antibodies against human α-synuclein and ubiquitin (**d**). The number of *h*α-Syn/ubiquitin-positive puncta was quantified (**e**). *p* values were calculated by Two-way ANOVA with Bonferroni Post Hoc Test. Scale bar, 10 µm. **(f-h)** The number of p62 puncta was quantified in microglia at striatum of brains either from AAV-*h*α-Syn-injected mice (n=3, **f, g**) or from *h*α-Syn-Tg mice (n=5, **h**) compared to AAV-GFP-injected mice (n=3) and non-Tg mice (n=3), respectively. *p* values were calculated by Mann–Whitney U test. Scale bar, 10 µm. **(i)** At 6-weeks post-AAV-*h*α-Syn administration into *Cx3cr1*^CreER-IRES-Eyfp^ mice, brain slices were stained with antibodies against GFP/YFP, human α-synuclein, p62 (left panel), and ubiquitin (right panel). Scale bar, 10 µm; 1µm in the magnified box. **(j)** Brain sections from 10-months-old *h*α-syn-Tg mice were fixed and stained using antibodies against human α-synuclein, Iba-1, and p62. Scale bar, 10 µm; 1µm in the magnified box. **(k, l, m)** Human α-synuclein were immunoprecipitated from AAV**-***h*α-Syn-injected brain (**k**), *h*α-Syn-Tg brain (**l**), and cultured microglia treated with 250nM *h*α-Syn protein for 6 hours (**m**). Arrows indicate the high-molecular-weight of p62. Data are representative of at least three independent experiments. All values are reported as mean ± SEM.

We next treated cultured microglia from *p62*-KO mice with *h*α-Syn (Fig. 3c). Remarkably, *h*α-Syn/ubiquitin-positive puncta were abolished in *p62*-KO cells (Fig. 3d, e); meanwhile, *p62* deficiency did not cause cell death regardless of *h*α-Syn treatment (Supplementary Fig. 4f), suggesting that p62 is necessary for the formation of *h*α-Syn/ubiquitin-positive puncta after *h*α-Syn ingestion, consistent with its role in selective autophagy^34^. In agreement with the *in vitro* results (Fig. 3a, b), the number of p62 puncta in microglia was significantly increased in the striatum of AAV9-*h*α-Syn injected mice at 6-weeks post-injection (Fig. 3f, g) and in 10-months-old *h*α-Syn-Tg mice compared to controls (Fig. 3h). Furthermore, p62/ubiquitin and *h*α-Syn puncta colocalized in EYFP-positive (AAV9-*h*α-Syn-injected *Cx3cr1*^CreER-IRES-Eyfp^) or Iba1-positive (*h*α-Syn-Tg mice) microglia (Fig. 3i, j). Through co-immunoprecipitation analysis, we found that *h*α-Syn interacted with specific p62 protein species of high-molecular-weight in AAV9-*h*α-Syn-injected brain, *h*α-Syn-Tg brain, and primary microglia treated with *h*α-Syn (Fig 3. k-m), supporting that oligomeric p62 directly binds and recruits *h*α-Syn into autophagosomes. The above data thus demonstrates that *h*α-Syn-induced p62 upregulation in microglia confers a selective autophagic degradation of ingested *h*α-Syn.

### α-synuclein-induced p62 upregulation requires TLR4 independently of TLR4 endocytosis

We next asked how microglia upregulate *p62* transcription in response to extracellular *h*α-Syn. Previous reports showed α-synuclein can trigger TLR4 or TLR2 activation in cultured glia, which leads to an increase in inflammatory gene transcription^40–42^. To test TLR4 involvement, we treated microglia from *Tlr4*-KO mice with *h*α-Syn and observed no increase in p62 protein and mRNA regardless of the dose (Fig. 4a, b). Also, the number of *h*α-Syn/ubiquitin-positive puncta present at 6 hours after *h*α-Syn treatment was significantly decreased in *Tlr4*-KO microglia compared to control cells (Fig. 4c). In addition, we treated cultures with TAK-242, an inhibitor of TLR4 signaling by selectively binding to Cys747 of TLR4 cytoplasmic tail that disrupts its interaction with adaptor molecules TIRAP and TRAM^43^ (Supplementary Fig. 5c). Similar to *Tlr4*-KO microglia, TAK-242-treated cells showed impaired p62 induction (Fig. 4d) as well as a reduction in the number of *h*α-Syn/ubiquitin-positive puncta following *h*α-Syn treatment (Fig. 4e). Furthermore, *Tlr4*-KO mice injected with AAV-*h*α-Syn showed reduced p62 puncta number in Iba1-positive microglia compared to control mice (Fig. 4f, g), indicating that TLR4 is required for microglial p62 induction in response to *h*α-Syn exposure both *in vivo* and *in vitro*.

**Figure 4.**
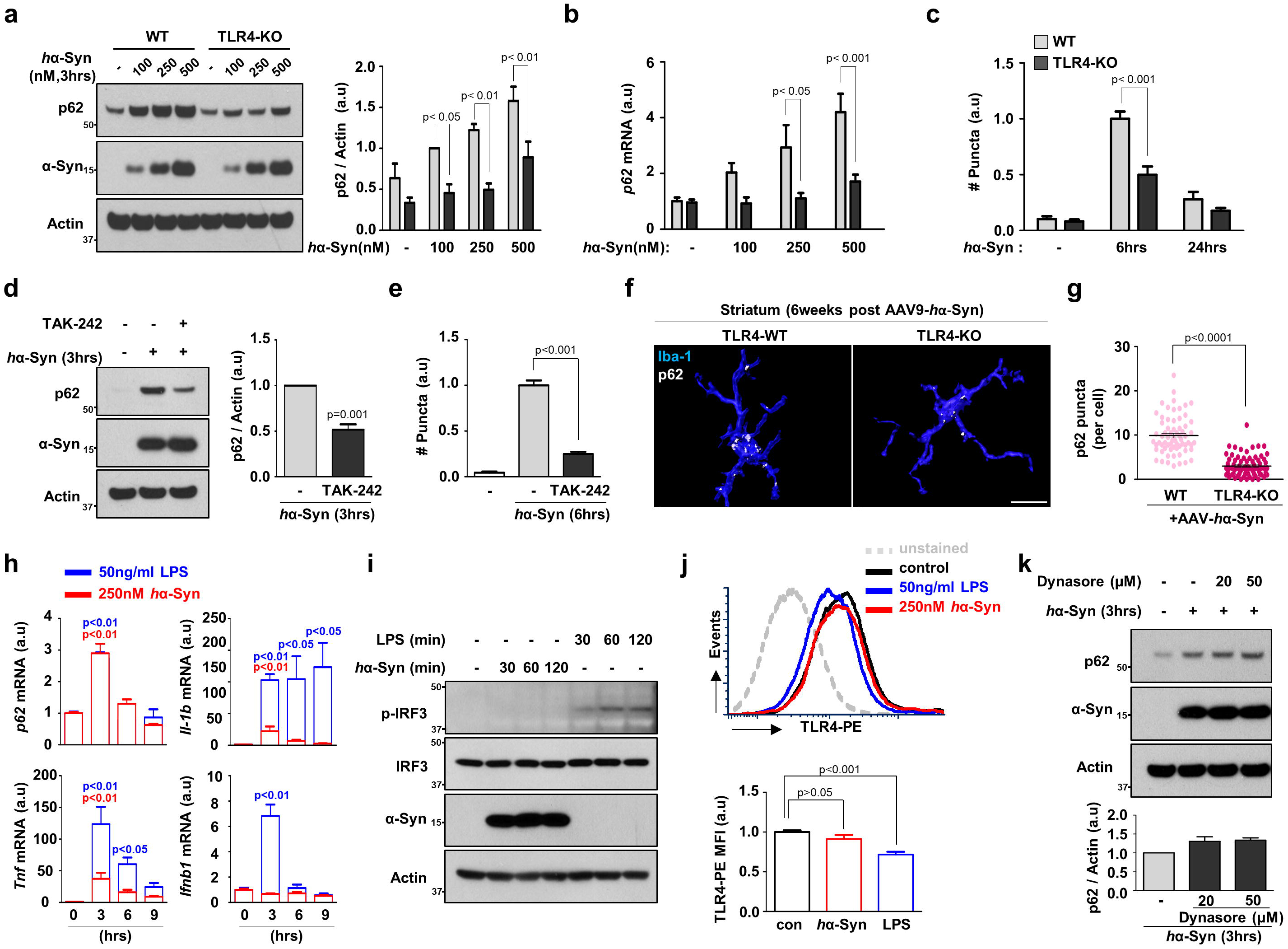
α-synuclein-induced p62 upregulation requires TLR4 independently of TLR4 endocytosis. **(a, b, c)** Microglia obtained from WT mice and *Tlr4*-KO mice were treated with *h*α-Syn protein and assayed for W.B (**a**), RT-qPCR (**b**) or for immunostaining (**c**). Band intensities were quantified in the right panel of a. The number of *h*α-Syn/ubiquitin-positive puncta was quantified (c). *p* values were calculated by Two-way ANOVA with Bonferroni Post Hoc Test. **(d, e)** Cells were pretreated with TAK242, a TLR4 signaling blocker, and treated with 250nM *h*α-Syn protein. The protein levels of p62 and the number of *h*α-Syn/ubiquitin-positive puncta were determined by W.B (**d**) and immunostaining (**e**), respectively. *p* values were calculated by Unpaired two-tailed Student’s t-test (d) and One-way ANOVA with Newman–Keuls post hoc test (e). **(f, g)** Representative 3D reconstruction pictures of microglia containing p62 in WT and *Tlr4*-KO mice injected with AAV-*h*α-Syn brains in (**f**). The number of p62 puncta was quantified in microglia at striatum of WT mice (n=5) and *Tlr4*-KO mice (n=5) (**g**). *p* values were calculated by Mann–Whitney U test. Scale bar, 10 µm. **(h)** Microglia were treated with either 250nM *h*α-Syn protein (red bar) or 50ng/ml LPS (blue bar) for indicated time and assayed for RT-qPCR. Each mRNA level was normalized to *Actin* mRNA. *p* values were calculated by One-way ANOVA with Newman–Keuls post hoc test. **(i)** After treating 250nM *h*α-Syn protein or 50ng/ml LPS for indicated time, cells were assayed for W.B using antibodies against p-IRF3 (S396), IRF3, and α-synuclein. **(j)** After treating *h*α-Syn protein or LPS for 30 minutes to microglia, the level of TLR4 receptor endocytosis was determined by detecting surface TLR4 using flow cytometry. Mean fluorescence intensity of TLR4 receptor staining on the surface was quantified. *p* values were calculated by One-way ANOVA with Newman–Keuls post hoc test. **(k)** Microglia were pretreated with Dynasore for indicated concentration and treated with 250nM *h*α-Syn protein for 3 hours. Data are representative of at least three independent experiments. All values are reported as mean ± SEM.

Lipopolysaccharide (LPS) triggers intracellular signaling via TLR4 binding, leading to the activation of the NF-κB and MAPK pathways. TLR4-LPS binding induces endocytosis of the TLR4-LPS complex and subsequent expression of interferon-beta 1 (*Ifnb1*) through activation of interferon regulatory factor 3 (IRF3)^44, 45^. Therefore, we asked if *h*α-Syn triggers similar TLR4 downstream events. Through RT-qPCR analysis, we found that the major target genes of the NF-κB pathway, *Il-1b* and *Tnf,* were up-regulated by LPS or *h*α-Syn treatment (Fig. 4h). Noticeably, *p62* was also increased by LPS as reported (Fig. 4h)^46^. However, LPS, but not *h*α-Syn, induced *Ifnb1* expression (Fig. 4h) and increased levels of p-IFR3 (S396), a marker of IRF3 activation (Fig. 4i), indicating the absence of IRF3 signaling in *h*α-Syn-treated microglia.

To test TLR4 endocytosis, we treated microglia with LPS or *h*α-Syn protein for 30 minutes and stained with TLR4 antibody conjugated with phycoerythrin (PE) to monitor the levels of TLR4 at the cell surface. LPS decreased the level of TLR4 on the plasma membrane as a result of endocytosis as reported^45^; in contrast, *h*α-Syn had little effect on TLR4 levels at the cell surface (Fig. 4j), suggesting that *h*α-Syn does not induce TLR4 internalization. Lastly, pretreatment with Dynasore did not block p62 induction by *h*α-Syn (Fig. 4k). The above observations show that the *h*α-Syn-TLR4 interaction upregulates p62 expression without causing TLR4 receptor endocytosis distinctively from LPS-TLR4 signaling.

### NF-**κ**B mediates α-synuclein**-**TLR4 signaling and p62 induction in microglia

Upon *h*α-Syn treatment, microglia activate NF-κB and p38 pathways as indicated by an increase of p-NF-κB (S536), a decrease of IκB level, and an increase of p-p38 (T180/Y182) (Fig. 5a). However, p-ERK1/2 (T202/Y204) and p-JNK (T183/Y185) were not changed (Fig. 5a). We next treated microglia with ML-120B, an inhibitor of IKK2, which is responsible for IκB phosphorylation and degradation that leads to NF-κB activation, and SB202190 or SB203580, two different inhibitors of p38. While the inhibitory effects of ML-120B, SB202190 and SB203580 were all confirmed in LPS-induced NF-κB activation and gene expression of proinflammatory cytokines (Supplementary Fig. 5d, e), only ML-120B significantly suppressed p62 induction and reduced the number of *h*α-Syn/ubiquitin-positive puncta after *h*α-Syn treatment (Fig. 5b, c), suggesting that NF-κB is the primary mediator for TLR4-triggered p62 induction. Supporting this notion, mRNA levels of *Il-1b* and *Tnf* were significantly lower in *Tlr4*-KO than WT cells, despite that both WT and *Tlr4*-KO microglia show an increase in *Il-1b* and *Tnf* expression after *h*α-Syn treatment (Fig. 5d). In contrast, NRF2 or TFEB pathways were unaltered as shown by the lack of change in the level of NQO-1 and HO-1, two major targets of NRF2, or the intensity of nuclear TFEB in response to *h*α-Syn^47, 48^ (Supplementary Fig. 4i, j).

**Figure 5.**
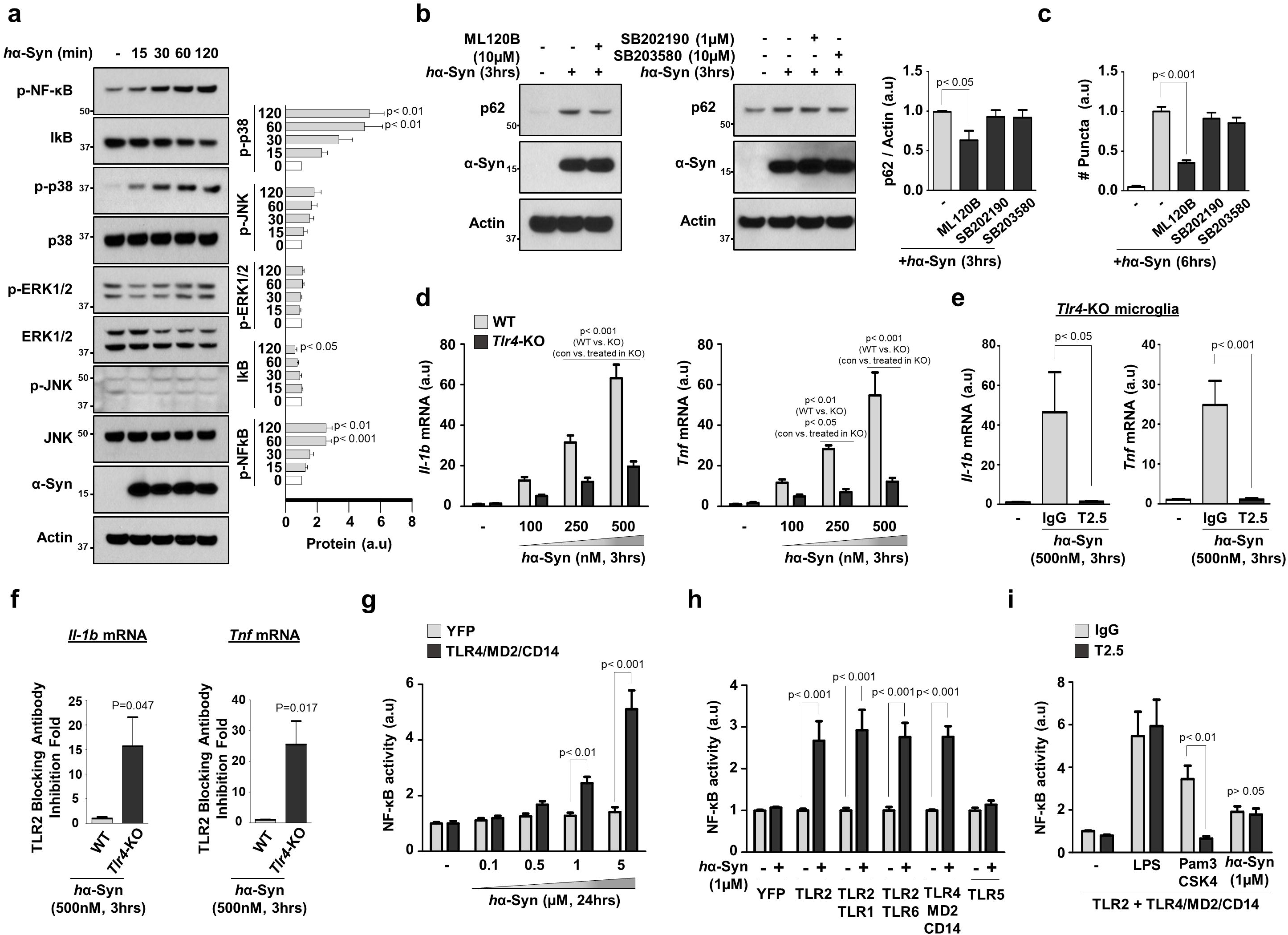
NF-κB mediates α-synuclein**-**TLR4 signaling and p62 induction in microglia. **(a)** WT microglia were treated with 250nM *h*α-Syn protein for the indicated time and assayed for W.B using antibodies against p-NF-κB (S536), IkB, p-p38 (T180/Y182), p38, p-ERK1/2 (T202/Y204), ERK1/2, p-JNK (T183/Y185), JNK, and human α-synuclein. The levels of proteins were quantified in the right panel. *p* values were calculated by One-way ANOVA with Newman–Keuls post hoc test. **(b, c)** Cells were pretreated with ML120B, an IKK-2 inhibitor, or SB203580 and SB202190, two different p38 inhibitors, for indicated concentrations before 250nM *h*α-Syn protein treatment, and assayed for W.B (**b**) or immunostaining (**c**). The levels of proteins were quantified in the right panel. The number of *h*α-Syn/ubiquitin-positive puncta was counted and quantified (c). *p* values were calculated by One-way ANOVA with Newman–Keuls post hoc test. **(d-f)** Microglia obtained from WT mice and *Tlr4*-KO mice were treated with *h*α-Syn protein for 3 hours with indicated concentration without (**d**) or with (**e, f**) TLR2-blocking antibody (T2.5), and assayed for RT-qPCR using primers for *Il-1b* (left panel) and *Tnf* (right panel). *p* values were calculated by Two-way ANOVA with Bonferroni Post Hoc Test (wt vs ko in d), One-way ANOVA with Newman–Keuls post hoc test (non-treated vs. treated in d, e), and Unpaired two-tailed Student’s t-test (f). **(g-i)** HEK293T cells were transfected with a luciferase vector that expresses luciferase under the promoter containing NF-κB-binding elements, and various TLRs. Then, cells were treated *h*α-Syn protein with indicated concentration (g, h, i), 50ng/ml LPS (i), or Pam_3_CSK_4_ (i) for 24 hours without (g, h) or with (i) 10µg/ml TLR2-blocking antibody (T2.5) and control IgG. *p* values were calculated by Two-way ANOVA with Bonferroni Post Hoc Test. Data are representative of at least three independent experiments. All values are reported as mean ± SEM.

### TLR4 is a preferred TLR for α-synuclein interaction to activate NF-**κ**B signaling

Previous studies suggest that in microglia α-synuclein interacts with TLR2^17^. We, therefore, tested whether the interaction between *h*α-Syn-TLR2 activates the NF-κB pathway. We applied a TLR2-blocking antibody (T2.5 clone), which suppressed p-NF-κB in response to the TLR2-specific ligand, Pam_3_CSK_4_ (Supplementary Fig. 5f). Remarkably, pretreatment with the TLR2-blocking antibody completely suppressed the induction of *Il-1b* and *Tnf* in response to *h*α-Syn in *Tlr4*-KO microglia (Fig. 5e). Interestingly, the TLR2-blocking antibody showed an enhanced effect of suppression in *Tlr4*-KO microglia compared to the WT (Fig. 5f), suggesting that TLR2 can partially compensate for TLR4 function in regulating the response to *h*α-Syn in *Tlr4*-KO microglia.

To test the specificity of *h*α-Syn effects on plasma membrane-expressed TLRs^49^, we established a NF-κB-luciferase system expressing a NF-κB response element and luciferase reporter gene in HEK293T cells expressing TLR2, TLR2/TLR1, TLR2/TLR6, TLR4/MD2/CD14 (TLR4 complex), or TLR5. First, we validated the specificity of this system by applying specific ligands for each TLR. Application of LPS, Pam_3_CSK_4_, and recombinant Flagellin peptides selectively increased NF-κB activity in TLR4 complex, TLR2 and TLR2 combinations, and TLR5, respectively (Supplementary Fig. 6). Furthermore, *h*α-Syn treatment significantly increased NF-κB activity in a dose-dependent manner in TLR4/MD2/CD14-transfected cells compared to YFP transfected cells (Fig. 5g). We also observed that *h*α-Syn increased NF-κB activity in TLR2, TLR2/1, and TLR2/6-transfected cells comparable to the activity in the TLR4 complex, while no change of the NF-κB activity was observed in YFP or TLR5-transfected cells (Fig. 5h). Furthermore, we applied the TLR2-blocking antibody in cells co-transfected with TLR2 and the TLR4 complex. As expected, pretreatment with TLR2-blocking antibody prevented the increase of NF-κB activity induced by Pam_3_CSK_4_ but not by LPS (Fig. 5i). Interestingly, application of the TLR2-blocking antibody had little effect on the increase of NF-κB activity induced by *h*α-Syn (Fig. 5i). These results suggest that *h*α-Syn-TLR4 interaction has a dominant effect over that of *h*α-Syn-TLR2 in regulating NF-κB signaling, while *h*α-Syn-TLR2 interaction only becomes important in NF-κB signaling in the absence of TLR4.

### Disruption of autophagy in microglia promotes accumulation of misfolded α-synuclein species in mouse brain

We next established microglia-specific *Atg7*-deficient mice (*Cx3cr1*^CreER-IRES-Eyfp^; *Atg7*^flox/flox^) by breeding *Cx3cr1*^CreER-IRES-Eyfp^ (tamoxifen-inducible Cre) with *Atg7*^flox/flox^ mice. Although previous studies showed that deletion of *Atg7* specifically in neurons causes neurodegeneration^50, 51^, we found that *Atg7*-deficiency had little effect on microglia number and state-specific gene signatures (Supplementary Fig. 7a-g), indicating that loss of autophagy does not cause microglial death or activation for at least up to 7-months following *Atg7* deletion.

We then tested the consequence of microglial *Atg7*-deficiency on α-synuclein homeostasis in the brain by investigating *Cx3cr1*^CreER-IRES-Eyfp^; *Atg7*^flox/flox^ mice expressing *h*α-Syn through either AAV injection (Fig. 6a) or transgenic expression (Fig. 6c). After injection of AAV9-*h*α-Syn, we observed that *Atg7*-deficient microglia contained more p62 puncta as well as p62/*h*α-Syn-positive puncta compared to control microglia (Supplementary Fig. 7h, i). Analysis of brain lysate fractions showed that microglial *Atg7*-deficient brains contained significantly elevated levels of detergent-insoluble *h*α-Syn at 6-weeks after AAV9-*h*α-Syn injection, as evidenced by a smear of high molecular weight *h*α-Syn species, compared to littermate controls (Fig. 6b).

**Figure 6.**
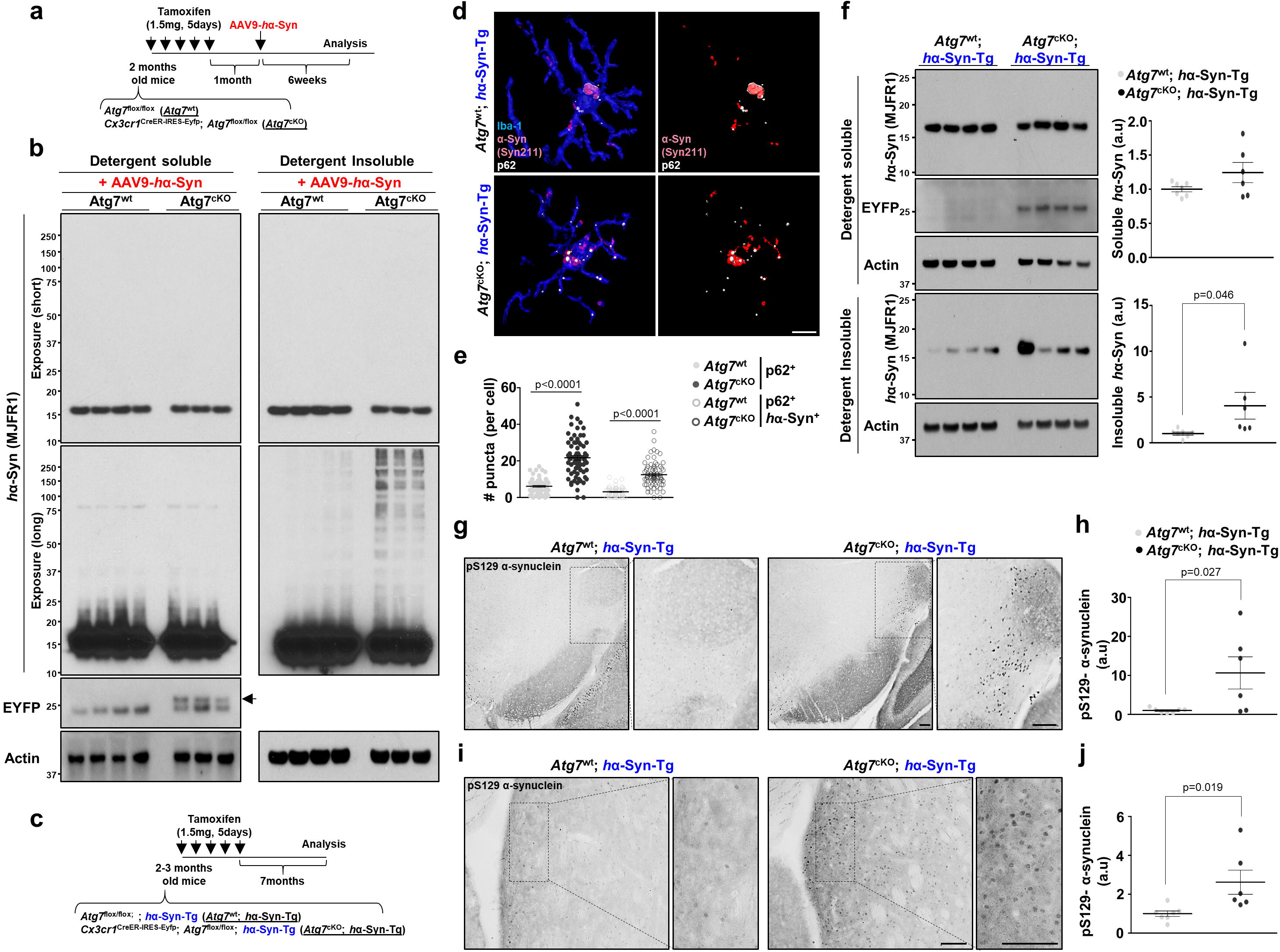
Disruption of autophagy in microglia promotes accumulation of misfolded α-synuclein species in mouse brain. **(a)** Experimental plan to test the roles of microglia-specific *Atg7*-deficiency in AAV-*h*α-Syn mice **(b)** After 6-weeks after AAV-*h*α-Syn, brains from *Atg7*^flox/flox^ mice (n=4) and *Cx3cr1*^CreER-IRES-Eyfp^; Atg7^flox/flox^ mice (n=3) were homogenized and fractionated into a detergent-soluble fraction and detergent-insoluble fraction and assayed for W.B using antibodies against human α-synuclein, and GFP/YFP. Actin were used as loading controls. Arrow indicates the EYFP band. **(c)** Experimental plan to test the roles of microglia-specific *Atg7*-deficiency in *h*α-Syn-Tg mice **(d, e)** After 7-months after tamoxifen, brain slices were fixed and stained with antibodies against human α-synuclein, p62, and Iba-1. Representative 3D reconstruction pictures of microglia containing p62 and human α-synuclein in the striatum of *Atg7*^flox/flox^ mice and *Cx3cr1*^CreER-IRES-Eyfp^; *Atg7*^flox/flox^ mice bred with *h*α-Syn-Tg mice were produced using Imaris (Bitplane, **d**). The number of p62 puncta or p62-positive/*h*α-Syn-positive puncta was quantified in microglia at striatum of *Atg7*^flox/flox^; *h*α-Syn-Tg mice (n=5) and *Cx3cr1*^CreER-IRES-Eyfp^; *Atg7*^flox/flox^; *h*α-Syn-Tg mice (n=5) (**e**). *p* values were calculated by Mann–Whitney U test. Scale bar, 10µm. **(f)** After 7-months after tamoxifen, brains from *Atg7*^flox/flox^; *h*α-Syn-Tg mice (n=7) and *Cx3cr1*^CreER-IRES-Eyfp^; *Atg7^f^*^lox/flox^; *h*α-Syn-Tg mice (n=6) were homogenized and fractionated into detergent-soluble fraction and detergent-insoluble fraction, and assayed for W.B using antibodies against human α-synuclein, and GFP/YFP for confirming the presence of *Cx3cr1*^CreER-IRES-Eyfp^. Band intensities were quantified in the lower panel. *p* values were calculated by Unpaired two-tailed Student’s t-test. **(g-j)** p-S129-α-synuclein-positive structures were visualized in brain sections from A*tg7*^flox/flox^; *h*α-Syn-Tg mice (n=7) and *Cx3cr1*^CreER-IRES-Eyfp^; *Atg7*^flox/flox^; *h*α-Syn-Tg mice (n=6) through DAB staining method (**g, i**), and quantified (**h, j**). *p* values were calculated by Unpaired two-tailed Student’s t-test. Scale bar, 100µm. All values are reported as mean ± SEM.

We also examined microglial *Atg7*-deficient mice after crossing with *h*α-Syn-Tg (*Cx3cr1*^CreER-IRES-Eyfp^; *Atg7*^flox/flox^; *h*α-Syn-Tg mice) at 7-months after *Atg7* deletion (Fig. 6c). The number of p62 puncta and p62/*h*α-Syn-positive puncta was significantly higher in *Atg7*-deficient microglia than control (Fig. 6d, e). Furthermore, microglial *Atg7*-deficient brains contained elevated levels of detergent-insoluble *h*α-Syn protein than control mice at 10-months of age (Fig. 6f). Furthermore, microglial *Atg7*-deficient mice showed increased staining of pS129 α-synuclein-positive structures particularly in the lateral SNpc (Fig. 6g, h) and the dorsal striatum (Fig. 6i, j).

### Loss of microglial *Atg7* promotes dopaminergic neuron degeneration associated with α-synuclein overexpression

Despite the lack of detectable neurodegeneration in *h*α-Syn-Tg mice or *Cx3cr1*^CreER-IRES-Eyfp^; *Atg7*^flox/flox^ mice (Supplementary Fig. 1f and Fig. 7c, d), the compound mice *Cx3cr1*^CreER-IRES-Eyfp^; *Atg7*^flox/flox^; *h*α-Syn-Tg (after the cross of the above two lines of mice) exhibited a decrease in the number of dopaminergic neurons as shown by stereological counting TH-positive neurons and Nissl-positive cells in the SNpc (Fig. 7a, b). These data suggest that disruption of microglial autophagy significantly enhances α-synuclein-mediated neurotoxicity. Taken together, our results demonstrate an important role for microglial autophagy in clearing α-synuclein released from neurons and preventing neurodegeneration.

**Figure 7.**
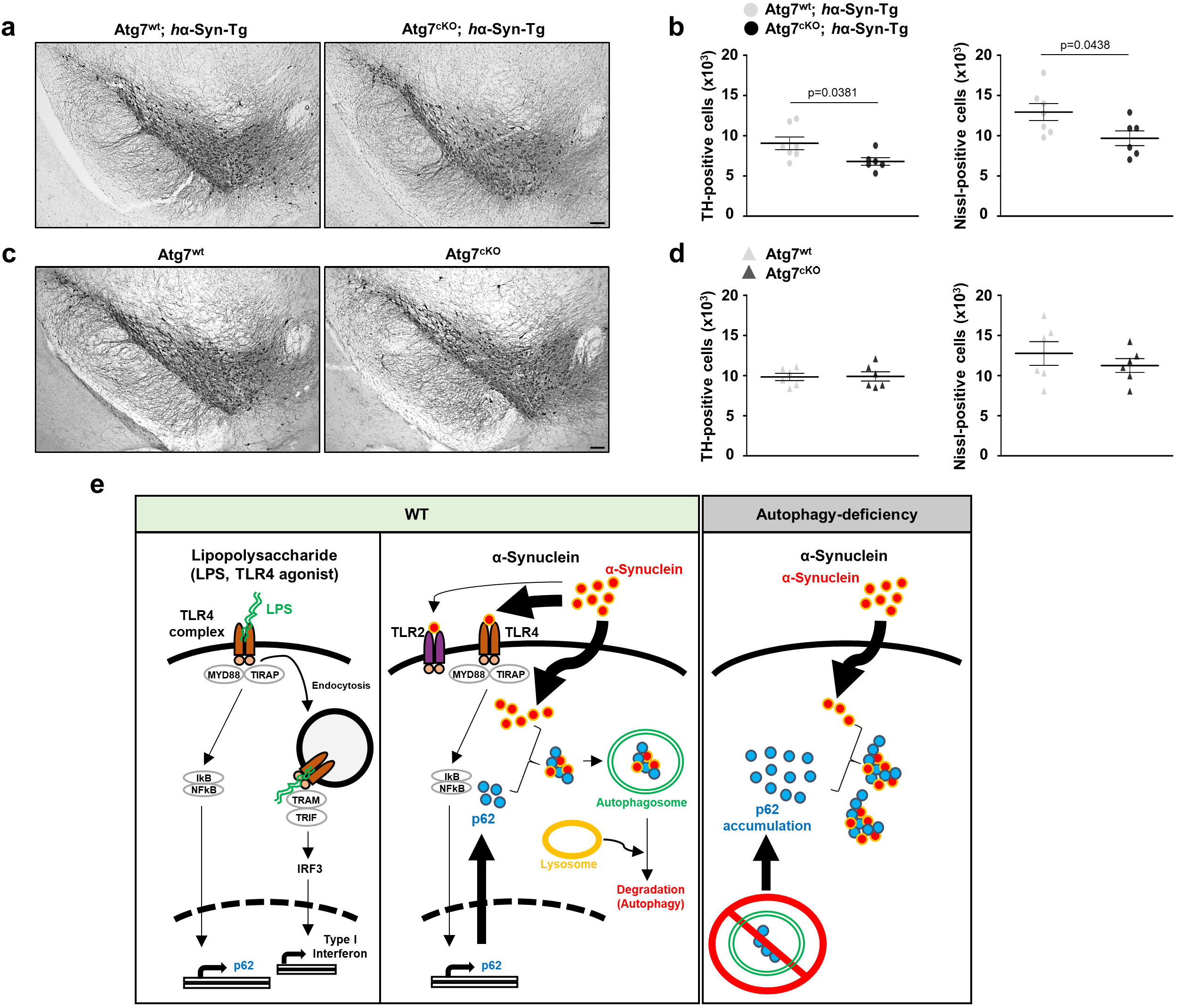
Loss of microglial *Atg7* promotes dopaminergic neuron degeneration associated with α-synuclein overexpression. **(a, b)** After 7-months after tamoxifen, brains from *Atg7*^flox/flox^; *h*α-Syn-Tg mice (n=7) and *Cx3cr1*^CreER-IRES-Eyfp^; *Atg7*^flox/flox^; *h*α-Syn-Tg mice (n=6) were fixed and stained using an antibody against tyrosine hydroxylase (TH) using DAB method (**a**). TH-positive cells and Nissl-positive cells in the SNpc area were counted (**b**). *p* values were calculated by Unpaired two-tailed Student’s t-test. Scale bar, 100µm. **(c, d)** After 12-months after tamoxifen, TH-positive cells and Nissl-positive cells were counted in SNpc area of *Atg7*^flox/flox^ (n=6) and *Cx3cr1*^CreER-IRES-Eyfp^; *Atg7*^flox/flox^ (n=6) brains. **(e)** Hypothetical model of microglial Synucleinphagy.

## Discussion

Our study has uncovered a significant role of microglia in clearing extracellular α-synuclein and protecting neurons through selective autophagy, which we term “synucleinphagy”. Here we show the first evidence that microglia sequester engulfed α-synuclein into autophagosomes for lysosomal degradation, thus solving a long-standing question as to whether wildtype α-synuclein is indeed degraded by autophagy. We identified the mechanism for microglia-specific synucleinphagy, which is mediated through TLR4-NF-κB signaling by transcriptional upregulation of the autophagy receptor, *p62/SQSTM1* (Fig. 7e). In contrast, neuron has no change of autophagosomes or p62 protein levels after the treatment with various forms of α-synuclein^52^ likely due to lack of TLR4-NF-κB pathway.

We demonstrate the critical role of TLR4-NF-κB-p62 in mediating synucleinphagy in microglia. TLR4 signaling has been linked to xenophagy, a selective autophagic degradation of infected bacteria, in macrophages. TLR4-NF-κB signaling was previously shown to induce selective autophagy after pathogen invasion^53–55^. In addition, p62 has a role in the clearance of intra-cytosolic *Shigella* and *Listeria*^56^. Thus, synucleinphagy and xenophagy have shared mechanism through TLR4-mediated signaling. Our study, however, showed that synucleinphagy require specifically p62 as other autophagy receptors such as Ndp52, Optineurin, or Nbr1, are not upregulated upon TLR4-NF-κB activation. We also noted that during synucleinphagy LC3-II level was unaltered, despite the increase of p62 levels and co-localization of LC3 puncta with ingested α-synuclein in microglia, suggesting a lack of global upregulation of autophagy but rather a restricted, local selective autophagy. The specific sequestration of α-synuclein into autophagosomes is likely mediated by p62 binding of ubiquitin-linked α-synuclein protein^34^; future study should identify the specific E3 ligase of α-synuclein in microglia responsible for synucleinphagy. Moreover, synucleinphagy differs from aggrephagy, a selective type of autophagy which degrades intracellular protein aggregates without the involvement of cell surface receptors (e.g. TLR) or NF-κB signaling.

Our result reveals that α-synuclein has a strong preference for TLR4 in the presence of both TLR4 and TLR2 receptors. Our observations are in an agreement with previous reports showing that both receptors interact with α-synuclein (albeit different forms). For example, TLR4 was required for NF-κB activation, and increased inflammation and phagocytic activity in response to various forms of α-synuclein^41, 42, 57^. In contrast, TLR2 also plays a role in neurons in response to α-synuclein^17, 58, 59^ in addition to microglia. Future studies should investigate further the significance of other TLRs in mediating α-synuclein degradation as well as inflammation in microglia. Our study also reveals a distinct feature of α-synuclein-triggered TLR4 activation, which does not promote TLR4 endocytosis in contrast to the effect of LPS. The absence of endocytosed TLR4 may prevent *Ifnb1* induction or increase of p-IFR3 (S396) level, despite the activation of NF-κB signaling, thus avoiding excessive inflammation and neurotoxicity^60^.

Finally, emerging human genomic studies suggest the autophagy-lysosome system as a converging pathogenic pathway of PD^61^, and many PD-related genes such as *PINK1*, *PARKIN*, *GBA*, and *LRRK2*, were shown to play a role in autophagy-related pathways in glial cells^62^. Our finding of microglial synucleinphagy via selective autophagy that confers neuroprotection should have a significant implication in understanding the role of microglia in preventing synucleinopathies. Our study thus opens a new avenue of research in therapeutic development by targeting synucleinphagy for the treatment of PD and DLB.

## Acknowledgments

This work was supported by an NIH grant (RO1, NS060123 and P50NS094733). We thank the assistance of members in the core facilities for Flow Cytometry, Microscopy, and qPCR at the Icahn School of Medicine at Mount Sinai. We thank NeuroScience Associates (Micheal J. Fox Foundation) for the Thy1-SNCA animal information and experiment. We are grateful to all members of Yue laboratory at the Icahn School of Medicine at Mount Sinai.

## Author contributions

Z.Y. conceived the project, supervised the entire study, and wrote a paper; I.C. designed, performed, and analyzed most of the experiments and wrote the paper; Y.Z. helped for the stereotaxic injection and stereological counting; S.S. performed NF-κB-luciferase experiments; M.P. helped *in vitro* results and analysis; Q.W. helped for analyzing the data; K.P. helped for preparing mice and writing a paper.

## Method

### Animals

B6.B10ScN-*Tlr4*^lps-del^/JthJ mice (*Tlr4*-KO, #007227), B6J.B6N(Cg)-*Cx3cr1*^tm1.1^(cre)^Jung^/J mice (*Cx3cr1*^Cre^, #025524), and B6.129P2(Cg)-*Cx3cr1*^tm2.1(cre/ERT2)Litt^/WganJ mice (*Cx3cr1*^CreER-IRES-Eyfp^ in this study, #021160) were purchased from Jackson Laboratory (Bar Harbor, ME). *Atg7*^flox/flox^ mice and *p62*-KO mice were kindly gifted from Dr. Masaaki Komatsu (Tokyo, Japan)^38, 63^. *Atg14*^flox/flox^ mice were Dr. Herbert W.Virgin (Washington University School of Medicine, St. Louis, MO)^39, 64^. GFP–LC3 transgenic mice (C57BL/6J) was described previously^35^. For generating microglia-specific *Atg7*-deficient mice, *Atg7*^flox/flox^ mice were bred to *Cx3cr1*^CreER-IRES-Eyfp^ (#021160) mice. *Cx3cr1*^CreER-IRES-Eyfp^; *Atg7*^flox/flox^ mice and littermate *Atg7*^flox/flox^ mice were injected with tamoxifen (intraperitoneal injection, five continuous days, 1.5 mg per mouse) to induce Cre recombinase expression. *Cx3cr1*^CreER-IRES-Eyfp^ mice express Cre recombinase not only in microglia but also in myeloid cell populations in the blood which are replenished by Cx3cr1-negative progenitors, which takes 1 month. However, due to limited turn-over, microglia will continuously express Cre recombinase^33^. Therefore, we injected AAV-*h*α-Syn after 1-month post tamoxifen administration to avoid any possible involvement of *Atg7*-deficient blood-derived cells in our observation. For obtaining *Atg7* or *Atg14*-deficient cultured cells, *Atg7*^flox/flox^ mice or *Atg14*^flox/flox^ mice were bred to *Cx3cr1*^Cre^ mice (#025524). All animal procedures were approved by Icahn School of Medicine at Mount Sinai Animal Care and Use Committee (IACUC-2015-0046).

### Generation of Thy1-SNCA mice (C57BL/6N-Tg(Thy1-SNCA)15Mjff/J mice, Jax #017682)

The transgenic construct contained a 4.2 kb murine Thy-1 (Gene location on chromosome 9: 43,851,467-43,856,662) promoter fragment containing ∼ 1.6 kb of 5’flanking sequence plus the 5’ first UTR exon (93bp), the first intron (2480 bp), and the non-coding region of exon 2 before ATG. This was followed by a 423 bp ORF of human wild-type α-synuclein cDNA (GenBank Acc# NM_000345), and 221 bp of SV40 polyA sequence in a pGL3 vector backbone (Promega). The above strategy was according to the published paper^65^. The microinjection fragment was purified and quantified before the injections. Three injection sessions were performed into C57BL/6Ntac embryos. Founders were identified and mated to wild-type C57BL/6NTac females. Analysis of integration site indicate insertion occurred at Chr11:40456787-40495044 and resulted in a 38.4 Kb deletion^66^. Copy number analysis was also performed on tail clips of the mice using the following primers and probe combination (spanning exons 2 and 3 of hSNCA gene):

Forward Primer: 5’-AGGACTTTCAAAGGCCAAGG-3’

Probe (FAM-labeled): 5’-AGTTGTGGCTGCTGCTGAGAAAACC-3’

Reverse Primer: 5’-CACTTGCTCTTTGGTCTTCTCAG-3’

The DNA copy number was determined by a standard curve method where for the standard curve the calculated number of copies of (1, 5, 10, 20, 50 and 100 copies) of the 5.1 kb transgene DNA fragment was mixed with 5 ng of DNA prepared from the wild type C57BL/6NTac mice. The data were analyzed by plotting ΔCt values against the known copy number. A standard curve was run in parallel with the samples prepared from the mice. An average copy number was 1.98 +/− 0.81. Following copy number analysis, the line was sent to The Jackson Laboratory for colony establishment (C57BL/6N-Tg (Thy1-SNCA)15Mjff/J; Stock No. 017682).

### Adeno-associated virus (AAV)

AAV vectors encoding either GFP or human WT α-synuclein were kindly gifted from Dr. Jia-Yi Li (Lund University, Lund, Sweden). AAV9-GFP or AAV9-human WT α-synuclein were packaged using the service provided by Vigene company (Rockville, MD). For establishing an acute PD animal model, 2µl of 6×10^13^ GC/ml of either AAV-GFP or AAV-α-synuclein were injected into the SNpc area of mice (AP: −3.28mm, ML: - 1.5mm, DV: −4.1mm) using a stereotaxic apparatus (Kopf Instruments, Tujunga, CA).

### Cell culture

Primary microglia were obtained from mixed glia cultured from forebrain of pups from 1-3 days old C57/BL6J WT mice (Jackson Laboratory). Briefly, forebrains were isolated and homogenized into single-cell suspensions by triturating with fire-polished Pasteur pipettes in Dulbecco’s modified Eagle’s medium (DMEM, #11965-092, Gibco™, Thermo Fisher Scientific) containing penicillin/streptomycin (#15080-063, Gibco™), GlutaMAX™ (#35050061, Gibco™), and heat-inactivated 10% fetal bovine serum (FBS, #S11550, Atlanta Biologicals, Atlanta, GA). Homogenized tissues were plated at 75 cm^2^ T-flask (BD Bioscience, San Jose, CA, USA) and incubated for 2-weeks. Floating microglia were detached from flasks by mild shaking and then filtered through a 70µm cell strainer to remove cell clumps or debris. Microglia were plated onto culture dishes at an appropriate density. HEK 293T cell line was maintained in DMEM supplemented with penicillin/streptomycin and 10% FBS.

### Adult microglia isolation

For adult microglia isolation, we adopted the established protocol ^32, 67^. Briefly, whole brains were isolated from mice after whole-body intracardial perfusion with ice-cold D-PBS (Gibco™) and homogenized using 15ml-size Dounce homogenizer in 5ml ice-cold D-PBS with 10 strokes. Homogenized tissues were filtered through a 70µm cell strainer to remove any debris and centrifuged with 1000 x g for 10 minutes at 4□. Cell pellets were resuspended in 5 ml of 70% Percoll^®^ (P1644, Sigma-Aldrich) made in 10% FBS-containing DMEM and 7 ml of 40% Percoll^®^ made in D-PBS were laid on the top of it gently in a 15ml tube (Falcon™, CA). After centrifugation with 800 x g speed for 30 minutes in RT, top layer enriched with myelin was removed and the middle layer (∼10ml) was mixed with ice-cold D-PBS (∼40ml), and centrifuged with 1400 x g for 5 minutes at 4□. This microglia-enriched collection was further used for flow cytometry.

### Flow Cytometry

Microglia-enriched pellets were resuspended in 500 µl of FACS buffer (1% FBS in ice-cold D-PBS), and incubated with 10 µg/ml of anti-mouse CD16/CD32 (#14-0161-82, Invitrogen) for 15 minutes on the ice to block Fc-receptor on microglia followed by 5 µg/ml of CD45 Monoclonal Antibody (I3/2.3) conjugated with APC-Cyanine7 (#A15395, Invitrogen) and 2 µg/ml of CD11b Monoclonal Antibody (M1/70) conjugated with PerCP-Cyanine5.5 (#45-0112-82, Invitrogen) for 30 minutes on the ice. After washed, the microglia population (CD11b^High^/CD45^Intermediate^) were gated and collected as shown in Supplementary Fig. 2. In an initial validation step, microglia from *Cx3cr1*^CreER-IRES-Eyfp^ were stained with CD45 and CD11b antibodies and the ratio of CD11b^High^/CD45^Intermediate^ cell population among EYFP-positive cells, and vice versa was checked by setting each other’s cell population as a parent gate as shown in Supplementary Fig. 2.

### Quantitative RT–PCR

cDNA was synthesized from total RNA with AffinityScript™ Multi-Temp RT (Stratagene) with oligo dT18 as primer. For real-time PCR, PlatinumTaq DNA polymerase (Invitrogen) and a SYBR green (Molecular Probes) containing buffer were used. The real-time PCRs were performed using a thermocycler (ABI7900HT; Applied Biosystems). The PCR conditions used were: 95°C for 2 minutes, 40 cycles of 95°C for 15 seconds, 55°C for 15 seconds, and 72°C for 30 seconds using the following primer pairs (Eurofins MWG Operon):

*Actin*, 5’-AGGTGACAGCATTGCTTCTC-3’ (sense), 5’-GCTGCCTCAACACCTCAAC-3’ (antisense);

*Axl*, 5’-AGCACAGTCTGCAAACTCC-3’ (sense), 5’-CTACCTCTAGCTCCGTAGGTT-3’ (antisense);

*ApoE*, 5’-AACCGCTTCTGGGATTACCTG-3’ (sense), 5’-CTCTCCCTCGGCTAGGCAT-3’ (antisense);

*Cd9*, 5’-TAACTTCATCTTCTGGCTCGCT-3’ (sense), 5’-AAACCAACCAGCATCATGAGG-3’ (antisense);

*Clec7a*, 5’-ACTTCAGCACTCAAGACATCC-3’ (sense), 5’-TGGCTTCCTTTCTCTGATCC-3’ (antisense);

*Csf1r*, 5’-CCTCAAACGTGGAGACACCAA-3’ (sense), 5’-CGTGTGCCAACATCATTGCT-3’ (antisense);

*Cst7*, 5’-TTCAACAACTGCACAAATGACA-3’ (sense), 5’-GGCCTTTCACCACCTGTACCA-3’ (antisense);

*Cx3cr1*, 5’-CAACCCCTTTATCTACGCCTT-3’ (sense), 5’-GACCCATCTCCCTCGCTTG-3’ (antisense);

*HexB*, 5’-TACAAGAACCAGTAGCCGTCCT-3’ (sense), 5’-CTCTAAACCTCGTAACGCTCC-3’ (antisense);

*H2-Ab1*, 5’-GCCCTCAACCACCACAACAC-3’ (sense), 5’-AGTCCCCATTCCTAATAAGCTGT-3’ (antisense);

*Il-1b*, 5’-CCTCTGATGGGCAACCACTT-3’ (sense), 5’-TTCATCCCCCACACGTTGAC-3’ (antisense);

*Il-6*, 5’-ACAGAAGGAGTGGCTAAGGA-3’ (sense), 5’-CGCACTAGGTTTGCCGAGTA-3’ (antisense);

*Ifnb1*, 5’-AGATGTCCTCAACTGCTCTC-3’ (sense), 5’-AGATTCACTACCAGTCCCAG-3’ (antisense);

*Ifng*, 5’-TGGCAGGAGATGTCTACACT-3’ (sense), 5’-GAAGCACCAGGTGTCAAGTC-3’ (antisense);

*Itgax*, 5’-CTGCTGCCACCAACCCTTC-3’ (sense), 5’-AGCCATCAATCAGGAACACGA-3’ (antisense);

*Lpl*, 5’-ACAAGTTTTAGAGCAGGACCAT-3’ (sense), 5’-TTGCACAGCAGTTTACAAGCATC-3’ (antisense);

*Olfml3*, 5’-GACACAGAACCCAGCTTTGC-3’ (sense), 5’-GCTACAGTCCGTCACCATATCGT-3’ (antisense);

*p62/Sqstm1*, 5’-GAAGCTGCCCTATACCCACA-3’ (sense), 5’-CCCGATGTCGTAATTCTTGGTC-3’ (antisense);

*P2ry12*, 5’-TTTGCTGGGCTCATCACGAAC-3’ (sense), 5’-ACTGAAGTAACTTGGCACACC-3’ (antisense);

*Tgfbr1*, 5’-GATCCATCACTAGATCGCCCTT-3’ (sense), 5’-CCGACCTTTGCCAATGCTT-3’ (antisense);

*Tmem119*, 5’-CTGACATTCTGGCTGCTACC-3’ (sense), 5’-CACCCTTCACAGGCTTTGCTC-3’ (antisense);

*Tnf*, 5’-TCACTGGAGCCTCGAATGTC-3’ (sense), 5’-GTGAGGAAGGCTGTGCATTG-3’ (antisense);

*Trem2*, 5’-GTCCCAAGCCCTCAACACC-3’ (sense), 5’-TCCTCACCCAGCTGCCGACA-3’ (antisense).

The cycle threshold (Ct) for the gene transcript was normalized to the average Ct for transcripts of the housekeeping gene, Actin, amplified in each reaction. Relative quantitation of normalized transcript abundance was determined using the comparative Ct method (ΔΔCt).

### Immunostaining

For brain tissue staining, mice were anesthetized and transcardially perfused with ice-cold PBS, and brains were obtained. Then, brains were fixed overnight at 4□ in 4% paraformaldehyde. Fixed brains were stored at 4□ in a 30% sucrose solution until they sank. Series of coronal sections (30 µm) were obtained with a cryostat (Leica, Wetzlar, Germany). Coronal sections were incubated with blocking/permeabilization buffer (5% goat serum and 0.25% Triton X-100 in PBS) at RT for 30 minutes and incubated with primary antibodies against Aquaporin 4 (AB3594, Millipore), GFAP (#130300, Thermofisher Scientific), GFP/YFP (MA5-15256, Thermofisher Scientific), Iba-1 (019-19741, Wako, Japan), NeuN (MAB377, Chemicon), p62 (GP62-C, Progen, Germany), α-synuclein (clone MJFR1, ab138501, Abcam; clone syn211 conjugated with Alexa594 fluorescein, sc-12767 AF594, Santa Cruz Biotechnology), or ubiquitin (clone P4D1, sc-8017, Santa Cruz Biotechnology) at 4□ overnight followed by secondary antibodies conjugated with Alexa-fluorescein. For 3,3’-diaminobenzidine (DAB) staining, serial sections were rinsed three times with PBS, treated with 3% H_2_O_2_ for 3 min, and rinsed with PBS containing 0.1% Triton X-100 (PBST). After incubating in blocking solution (10% Goat serum and 0.25% Triton X-100 in PBS), sections were incubated overnight at room temperature with primary antibodies against Tyrosine hydroxylase (AB152, Millipore) and pS129 α-synuclein (ab51253, Abcam). After rinsing in PBST, sections were incubated with biotinylated secondary antibodies (Vector Laboratories, Burlingame, CA, USA) for 1 hour and the avidin/biotin system (Vector Laboratories) for 30 minutes and visualized using a DAB solution (Vector Laboratories). Sections were then mounted on gelatin-coated slides and examined under a bright-field microscope (Olympus Optical, BX51, Tokyo, Japan). For cultured primary microglia, cells were fixed with 4% paraformaldehyde at 4□ for 15 minutes and incubated with microwave-boiled antigen retrieval solution (#CTS013, R&D System) for 5 min. After washing with PBS, cells were incubated with blocking/permeabilization buffer (1% BSA and 0.1%Triton X-100 in PBS) at RT for 30 minutes and incubated with primary antibodies including EEA1 (610457, BD Bioscience), α-synuclein (clone MJFR1, ab138501, Abcam; clone 2f12, MABN1817, Millipore; clone 42, 610787 BD Bioscience; clone 204, #2647, Cell Signaling), p62 (GP62-C, Progen, Germany), ubiquitin (clone P4D1, sc-8017, Santa Cruz Biotechnology), and TFEB (A303-673A, Bethyl Laboratories Inc, TX) at 4□ overnight followed by secondary antibodies. Images were obtained using a confocal microscope (Zeiss LSM780, Carl Zeiss, Jena, Germany).

### Microglia cell number and morphology analysis

From two brain slices from each animal, z-stack pictures were obtained with 20x magnification at multiple areas of the striatum using a confocal microscope after staining with Iba-1 antibody (Zeiss LSM 780, Carl Zeiss, Jena, Germany). Then, microglial cell number was counted from z-stack projected pictures using ‘cell counter’ plug-in in Fiji software (National Institutes of Health, MD). For morphology analysis, the length of processes and terminal points of branches were measured by a semi-automated way using the ‘filament’ plug-in in Imaris 3D interactive visualization software (Bitplane, Zurich, Switzerland). To show the colocalization of two interest proteins or the location of proteins in microglia, any immunostaining signals outside of Iba-1-staining area were masked using ‘surface’ and ‘mask’ plug-in in Imaris software and each protein was made into the surface according to each staining signal. Then, final 3D rendering surface images were made by combining each protein surface to make a representative image.

### Brain Fractionation

Frozen brains (one hemisphere each animal) were homogenized with Dounce homogenizer in 2 ml of sucrose buffer (0.32M Sucrose in 50mM Tris-HCl, pH7.4) added with Halt™ Protease and Phosphatase Inhibitor Cocktail (#78440, Thermofisher Scientific), and spun-down for 10 min at 1000 x g 4□. The supernatant (∼1.3ml) was collected and 0.9ml of lysates were mixed with 0.1ml of 10% Triton X-100 to make 1%Triton X-100 in sucrose buffer. After spin-down for 10 min at 16000 x g 4□, the supernatant was collected into the new tube (detergent-soluble fraction), and the pellet was washed with 1%Triton X-100 two times to remove any soluble proteins. Finally, the washed pellet was resuspended in 0.1ml of 1%SDS/1%Triton X-100/Sucrose buffer and sonicated with the probe-tip sonicator in level ‘2’ for 10 sec (Fisher Scientific 550 Sonic Dismembrator). After spin-down for 5min at 16000 x g RT, the supernatant was collected into the new tube (detergent-insoluble fraction). Protein concentration was determined by Pierce BCA Protein Assay Kit (#23225, Thermofisher Scientific).

### Immunoprecipitation

Brain lysate from 1%Triton X-100/Sucrose buffer fractions or cultured microglia lysed in 1%Triton X-100 PBS were incubated with α-synuclein antibody (clone 2f12, MABN1817, Millipore) in 4□ overnight followed by Dynabeads™ Protein G (#10009D, Thermofisher Scientific). After washing with 1%Triton X-100 PBS three times, proteins that bound to beads were released by boiling in LDS sample buffer (Thermofisher Scientific) at 95□ for 10 min.

### Western blot analysis

Cultured cells were lysed on ice in RIPA buffer (50□mM Tris-HCl (pH 7.4), 1% NP-40, 1□mM NaF, 0.25% Na-deoxycholate, 1□mM Na_3_VO_4_ and 150□mM NaCl) containing Halt™ Protease and Phosphatase Inhibitor Cocktail (#78440, Thermofisher Scientific). For FACS-sorted microglia, cells were lysed in 1% Triton/8M Urea/PBS buffer to maximize the yield, sonicated with the probe-tip sonicator in level ‘2’ for 10 sec (Fisher Scientific 550 Sonic Dismembrator). For AAV model, total ∼30000 microglia sorted from FACS were used and, for transgenic mice, ∼120000 cells were used for each group. Lysates were centrifuged, and proteins in the supernatant were separated by NuPAGE® Precast Gel System (Thermofisher Scientific). Membranes were incubated with primary antibodies including Actin (#3700S, Cell Signaling), ATG7 (MAB6608, Cell Signaling), ATG14 (PD026, MBL), Dopamine transporter (MAB369, Millipore), ERK1/2 (#9107S, Cell Signaling), p-ERK1/2 (Thr202/Tyr204, #4370P, Cell Signaling), GFP/YFP (MA5-15256, Thermofisher Scientific), IκB (#4814, Cell Signaling), p-IRF3 (S396, #29047, Cell Signaling), IRF3 (#4302, Cell Signaling), JNK (#9252S, Cell Signaling), p-JNK (Thr183/Tyr185, #4671S, Cell Signaling), LC3A/B (#12741S, Cell Signaling), NBR1 (16004-1-AP, Proteintech), NDP52 (12229-1-AP, Proteintech), NQO-1 (11451-1-AP, Proteintech), HO-1 (sc-10789, Santa Cruz) p-NF-κB (S536, #3033S, Cell Signaling), Optineurin (10837-1-AP, Proteintech), p38 (#9212S, Cell Signaling), p-p38 (Thr180/Tyr182, #4511S, Cell Signaling), p62 (PM066, MBL), p62 (18420-1-AP, Proteintech), α-synuclein (clone MJFR1, ab138501, Abcam; clone 2f12, MABN1817, Millipore; clone 42, 610787 BD Bioscience; clone D37A6, # 4179, Cell Signaling), or Tyrosine hydroxylase (T2928, Sigma) at 4□ overnight, washed with Tris-buffered saline containing 0.1% Tween 20, incubated with secondary antibodies, and visualized with SuperSignal™ West Pico Chemiluminescent Substrate (Thermofisher Scientific).

### HEK293T-NF-κB luciferase assay system

HEK293T cells were seeded into a 96-well plate at a density of 2 × 10^4^ cells per well overnight. Cells were transfected using Lipofectamine® 2000 (Invitrogen) with pcDNA3 vectors encoding either TLR1 (Addgene #13014), TLR2 (Addgene #13015), TLR4 (Addgene #13018), MD2 (Addgene #13028), CD14 (Addgene #13645), TLR5 (Addgene #13019), or TLR6 (Addgene #13020) with pGL4.32[luc2P/NF-κB-RE/Hygro] vector containing five copies of an NF-κB response element (NF-κB-RE) that drives transcription of the luciferase reporter gene luc2P (*Photinus pyralis*) (#N111A, Promega, Madison, WI). Vectors were gifts from the Doug Golenbock Lab through Addgene. After 24 hours, cells were treated with either EndoClear□□ human recombinant α-synuclein (AS5555, Anaspec, CA), lipopolysaccharide (LPS, #L7770, Sigma-Aldrich), Pam_3_CSK_4_ (#tlrl-pms, Invivogen, CA), or recombinant Flagellin (Rec FLA-ST, #tlrl-flic-10, Invivogen, CA). Following 24 hours of stimulation by the ligand, the luciferase activity was measured using the ONE-Glo™ Luciferase assay system (#E6110, Promega, Madison, WI).

### Electron Microscope

Cell cultures grown on Permanox slides (Electron Microscopy Sciences (EMS), Hatfield, PA) were taken from incubation, and directly placed in 2% glutaraldehyde (EMS) and 2% paraformaldehyde (EMS)/ 0.1M Sodium Cacodylate buffer (EMS) for a minimum of 2 hours at 4□. Cells were washed, fixed in 1% aqueous osmium tetroxide at RT for 1 hour, washed, and transferred to 2% aqueous uranyl acetate at RT for 1 hour. Slides were washed with distilled water, dehydrated in an ascending aqueous ethanol series, and then embedded in Epon resin (EMS). Inverted BEEM capsules (#3, EMS) were placed directly over regions of interest, filled with fresh resin, and transferred to a vacuum oven for heat polymerization at 60□ for 12-24 hours. To separate the cells from the slides, a hot plate was heated to 60°C, and the slides placed directly on a pre-heated hot plate for exactly 3 minutes and 30 seconds. The capsules were removed from the hot plate and the capsules carefully dislodged free from the slide using a plier. Ultrathin (85 nm) sections were collected onto 300 mesh copper grids (EMS) using a Leica UC7 ultramicrotome (Leica Biosystems Inc., Buffalo Grove, IL), contrast stained with uranyl acetate and lead citrate, and imaged on a Hitachi 7700 transmission electron microscope (Hitachi High Technologies America, Inc., Dallas, TX) equipped with an AMT 2K x 2K digital camera (Advanced Microscopy Techniques, Corp., Woburn, MA).

### Image Analysis and Quantification

For counting p62-positive or p62/α-synuclein-positive puncta in microglia *in vivo*, fixed brain slices were stained with primary antibodies against human α-synuclein, p62, and Iba-1 and z-stack pictures were obtained with 63x magnification (1024×1024 pixels) at multiple areas of the striatum using a confocal microscope (Zeiss LSM 780, Carl Zeiss). Puncta were counted inside of Iba-1 positive area in every single focal plane throughout the whole z-stack manually. In cultured cells treated with EndoClear□□ human α-synuclein (AS5555, Anaspec, CA), pictures were taken with 20x magnification from multiple areas (up to 15 areas per coverslip), and ubiquitin/α-synuclein-positive puncta were counted by a plug-in ‘Cell counter’ in Fiji software (National Institutes of Health, MD). Then, the number of puncta each picture was divided by cell number and normalized to each control group. For quantification of pS129 α-synuclein DAB staining, pictures were taken with 20x magnification and positive signals were counted by a semi-automated way by using ‘Find Maxima’ in Fiji software (National Institutes of Health, MD) after setting the threshold. Values were normalized to the average values from the control group in each cohort. For stereology counting of TH-positive or Nissl-stained cell bodies in the SNpc, 1 in every 6 sections was selected with a random start and a total of 5 brain slices on average were used for each mouse for IHC labeling for TH, including DAB enhancement, followed by cresyl violet staining to reveal all neurons. A Zeiss Axioplan2 was used for tissue slice imaging with a 20× objective, and Stereo Investigator was used to estimating the total number of neurons in the region of interest using the following parameters: frame sizes, 150 × 150 μm; grid sizes, 250 × 250 μm; top guard zone height, 2 μm; and optical dissector height, 8 μm. These parameters yielded a coefficient of error <10% throughout the analysis.

### Statistics and Reproducibility

The statistical significance of differences between two groups was determined using the unpaired two-tailed Student’s t-test or Mann–Whitney U test based on the normality test. For multiple-means comparisons, statistical significance was determined by one-way analysis of variance followed by Newman–Keuls post hoc test or two-way analysis of variance with Bonferroni post hoc test using Graph Pad Prism 5 (GraphPad Software, CA, USA). All values are reported as mean ± SEM. Data are representative of at least three independent experiments unless indicated.

