## Supplemental Figure for "Microglia Clear Neuron-released α-Synuclein via Selective Autophagy and Prevent Neurodegeneration"

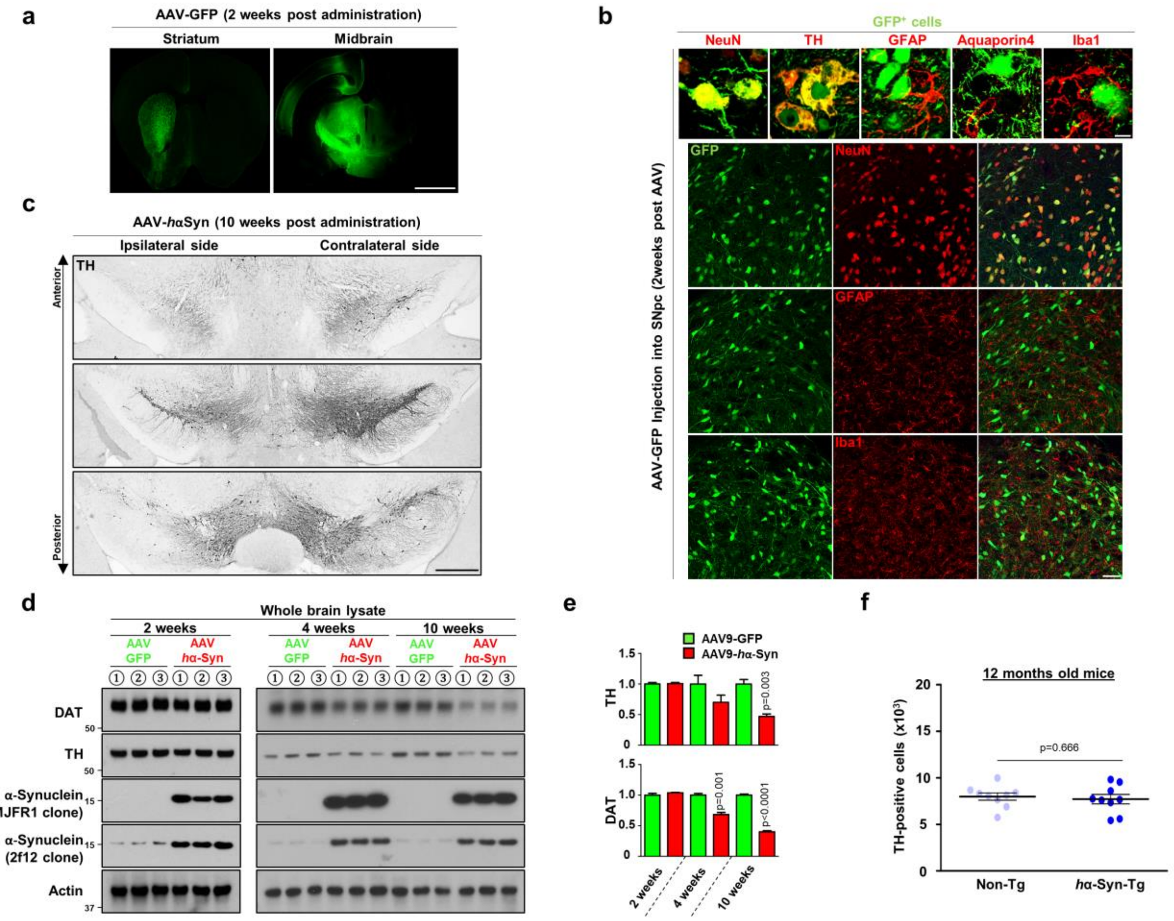


**Supplementary Figure 1. Characterization of two different PD model overexpressing human α-synuclein.**

**(a)** AAV9 expressing GFP under the neuron-specific synapsin promoter (AAV-GFP) was administered into substantia nigra and GFP expression was examined after 2-weeks. Scale bar, 2mm.

**(b)** To confirm the neuron-specific expression of GFP, brain slices were stained with antibodies against neuronal proteins (NeuN and Tyrosine hydroxylase (TH)), astrocytes markers (GFAP and Aquaporin 4), and microglia marker (Iba-1). Scale bar, 5 μm in the upper panel, 50 μm in the lower panel.

**(c)** Loss of dopaminergic neurons in the ipsilateral side of SNpc was visualized by tyrosine hydroxylase (TH) staining with DAB staining method in three different coronal sections at 10-weeks post-AAV-*h*αSyn administration. Scale bar, 500 μm.

**(d, e)** After AAV-GFP or AAV-*h*α-Syn administration, whole brains were homogenized and assayed for W.B using antibodies against DAT, TH, and α-synuclein at indicated time (d). Actin was used as loading controls. The intensities of protein bands were quantified (e). *p* values were calculated by Unpaired two-tailed Student's t-test.

**(f)** Brains of 1-year-old non-Tg (n=9) and *h*α-Syn-Tg mice (n=9) aged mice per genotype were fixed and stained using an antibody against tyrosine hydroxylase (TH) using DAB method. $TH^{+}$-positive cells were counted.

All values are reported as mean ± SEM.

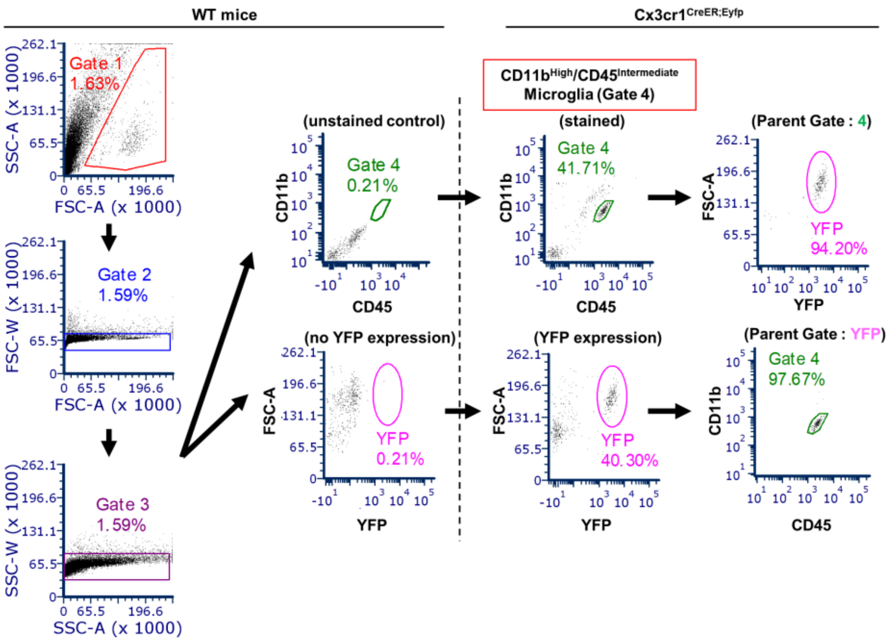


**Supplementary Figure 2. Gating strategy for obtaining CD45$^{intermediate}$ and CD11b$^{high}$ microglia from adult mouse brain.**

Enriched microglia population prepared from a Percoll®-gradient method were stained with CD45 and CD11b antibodies conjugated with APC-Cy7 and Percp-Cy5.5, respectively. Microglia cell population were gated at the first (Gate 1) and gated for single cell population (Gate 2 and 3). CD45$^{intermediate}$ and CD11b$^{high}$ cells (Gate 4 population) were collected. See the detail information in Method.

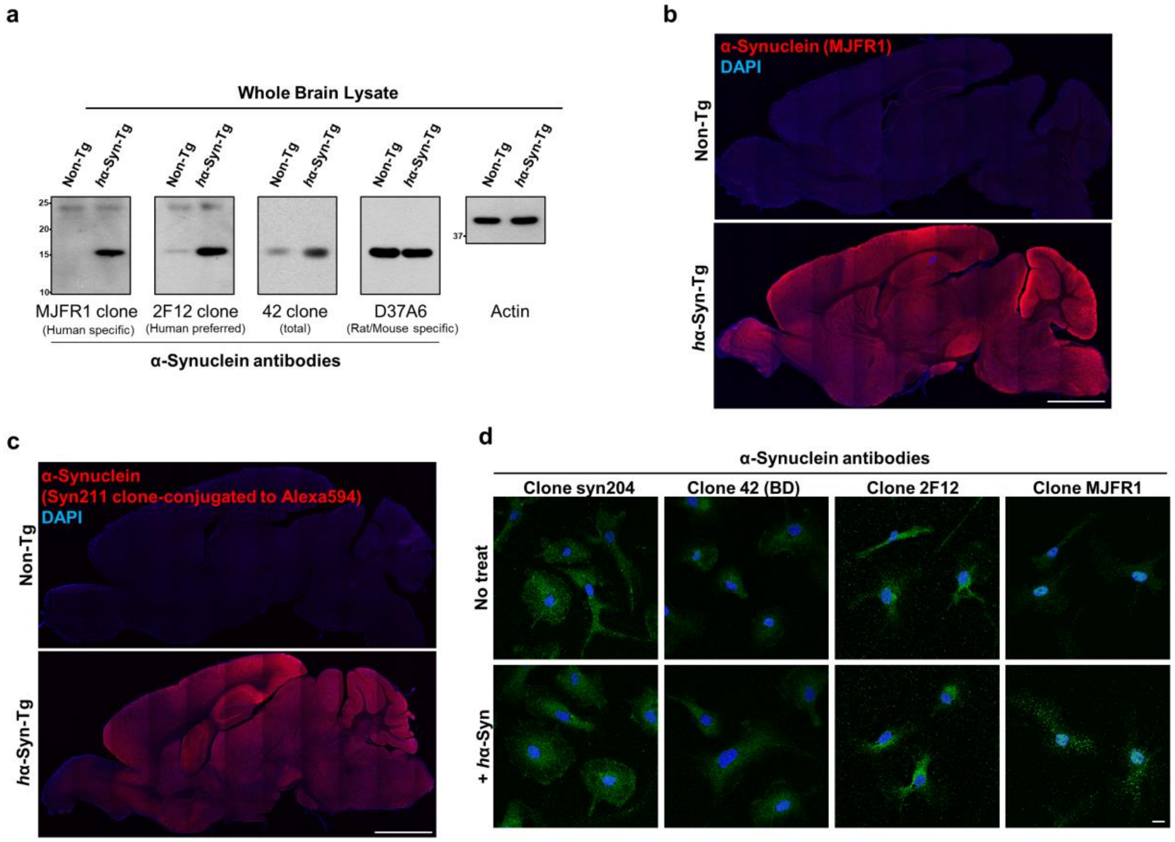


**Supplementary Figure 3. Test the specificity of various α-synuclein antibodies.**

**(a-c)** Brains from 3-months old non-Tg and *h*α-Syn-Tg mice were homogenized in TBS/Sucrose buffer and assayed for W.B with indicated antibodies against α-synuclein (a) or fixed and stained with human α-synuclein antibodies (MJFR1 clone in b; Syn211 clone in c). Actin was used as a loading control. Scale bar, 2mm.

**(d)** Cultured primary microglia were stained with indicated human α-synuclein antibodies in the presence/absence of 250nM recombinant human α-synuclein treatment for 1 hour followed by Alexa488-secondary antibody. Scale bar, 10μm.

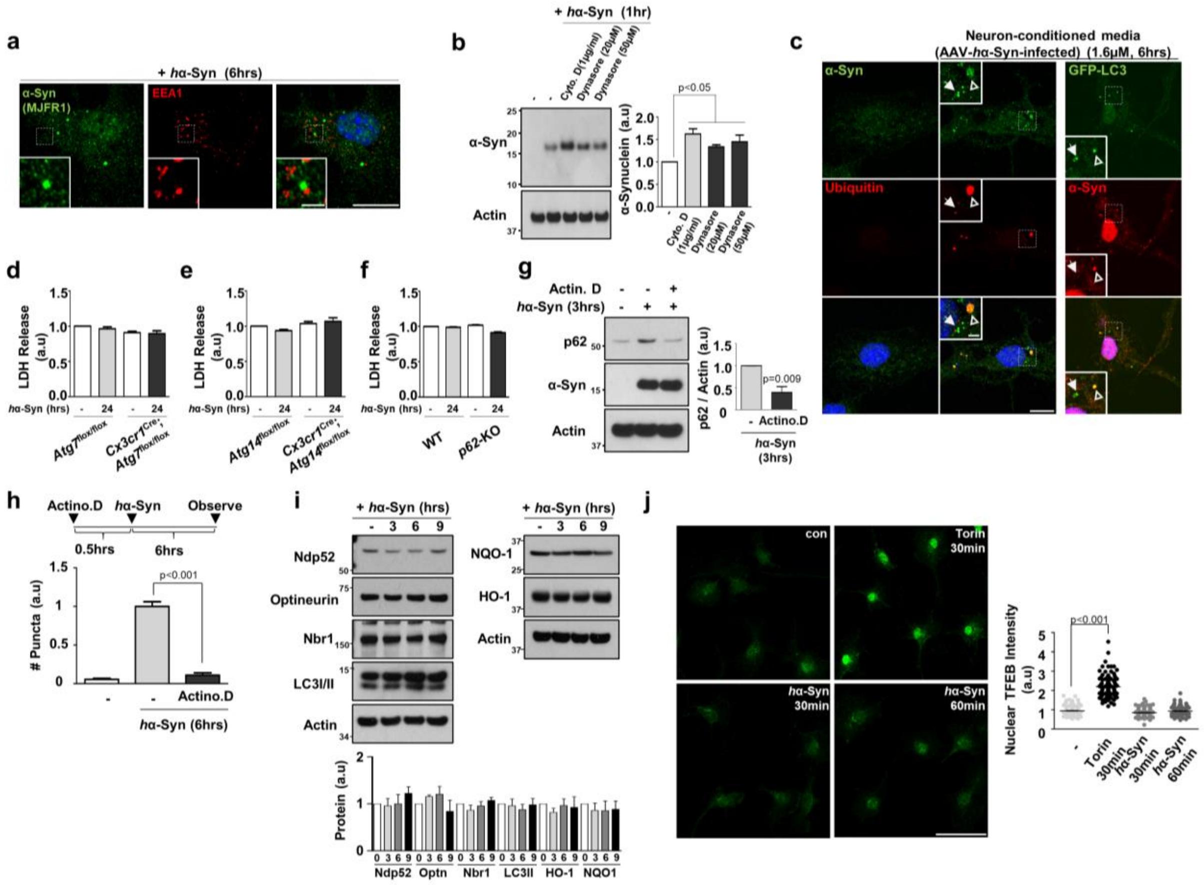


**Supplementary Figure 4. Characterization and effects of human α-synuclein on cultured microglia.**

**(a)** Microglia were treated 250nM *h*α-Syn protein for 6 hours and stained using antibodies against human α-synuclein (MJFR1 clone) and EEA1. Scale bar, 10 μm; 5 μm in magnified boxes.

**(b)** Microglia were pretreated with cytochalasin D (Cyto. D), an inhibitor of actin polymerization, and Dynasore, an inhibitor of endocytosis, before adding 250nM *h*α-Syn for 1 hour. Band intensities were quantified in the right panel. Acin was used a loading control. *P* values were calculated by One-way ANOVA with Newman–Keuls post hoc test.

**(c)** Embryonic mice cortical neuron was prepared and infected AAV9-α-*h*Syn (10µl of $6\times10^{13}$ GC/ml virus into 2ml medium) at DIV3 and the medium (containing ~3.2µM of *h*Syn) was collected at DIV14. Cultured wt microglia (left panel) and GFP-LC3 microglia (right panel) were treated with neuron-conditioned medium by exchanging the half of medium for 6 hours, and stained. Scale bar, 10µm; 2µm in the magnified box.

**(d-f)** The levels of lactate dehydrogenase (LDH) were measured using LDH-Cytotoxicity Colorimetric Assay Kit (#K311, Biovision, Mountain View, CA) according the manufacturer's instructions in the medium collected from cultured microglia either from *Atg7*$^{\text{flox/flox}}$ mice (**d**) or *Atg14*$^{\text{flox/flox}}$ mice (**e**) with or without *Cx3cr1*$^{\text{Cre}}$, and *p62*-KO mice (**f**) in the absence/presence of 250nM *h*α-Syn protein.

**(g, h)** Microglia were pretreated with Actinomycin D, a blocker of mRNA translation, for 30 minutes and treated with 250nM *h*α-Syn protein. Levels of p62 protein were quantified in the lower panel. The number of *h*α-Syn/ubiquitin-positive puncta was counted and quantified (g). *P* values were calculated by Unpaired two-tailed Student's t-test (**g**) and One-way ANOVA with Newman–Keuls post hoc test (**h**).

**(i)** After 250nM *h*α-Syn protein treatment, cells were lysed and assayed for W.B using antibodies against NDP52, Optineurin, NBR1, LC3 I/II, NQO-1, and HO-1. Actin was used as a loading control. Band intensities were quantified in the lower panel.

**(j)** After 250nM *h*α-Syn protein treatment or Torin, cells were fixed and stained using antibody against TFEB. Intensities of Nuclear TFEB were measured. Scale bar, 50µm.

All values are reported as mean ± SEM.

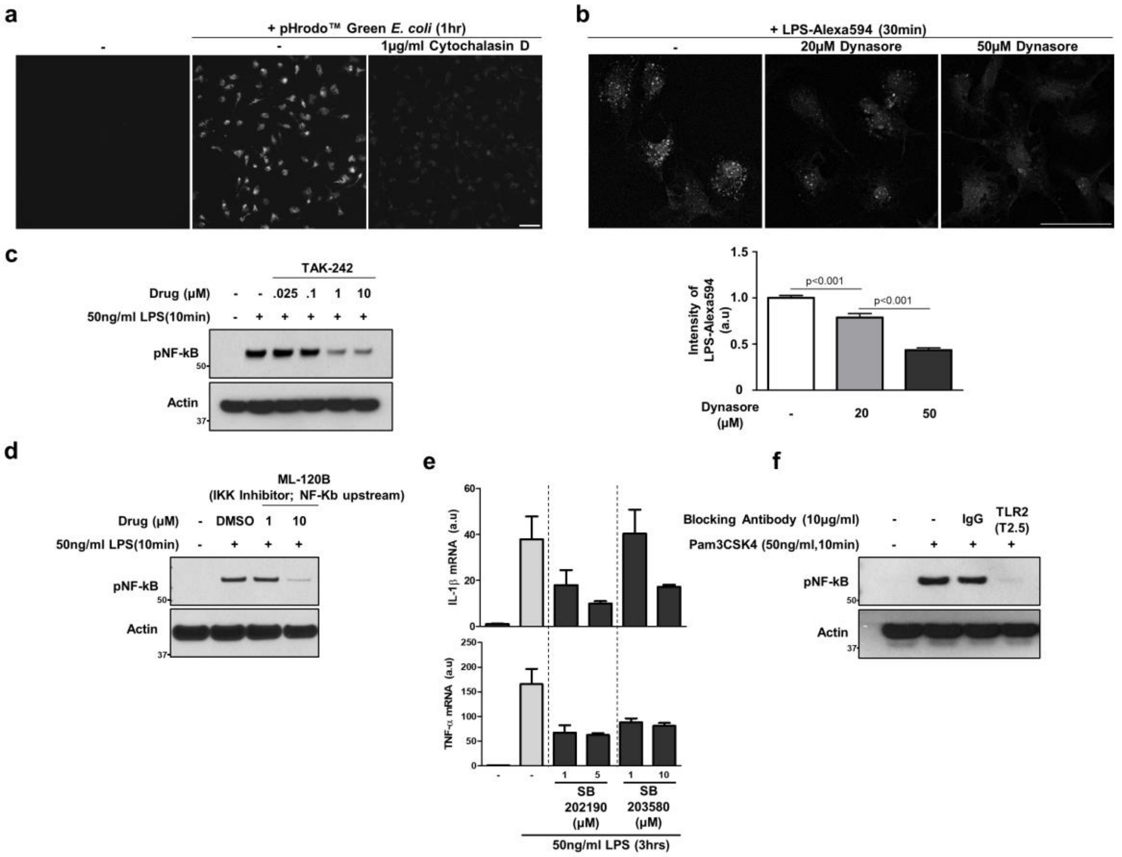


**Supplementary Figure 5. Drugs and TLR2 blocking antibody test.**

**(a, b)** Microglia were treated with Cytochalasin D (a) and Dynasore (b) as indicated concentrations before adding pHrodo™ Green *E.coli* (#P35366, Thermofisher Scientific) (a) and LPS conjugated with Alexa594 (L-23353, Thermofisher Scientific) (b), respectively, according the manufacturer's instructions. p values were calculated by One-way ANOVA with Newman–Keuls post hoc test (b). Scale bar, 50 µm

**(c-e)** Microglia were pretreated with TAK242 (a), ML-120B (b, an IKK2 inhibitor, #4899, Tocris), SB203580 and SB202190 (c, two different p38 inhibitors, #1202 and #1264, Tocris) before adding 50ng/ml LPS (L7770, Sigma), a TLR4-specific agonist, for indicated time. The protein levels of p-NF-κB (**c, d**) and mRNA levels of *Il-1b* and *Tnf* (**e**) were examined by W.B and RT-qPCR, respectively.

**(f)** Cells were pretreated with 10 μg/ml of TLR2 blocking antibody (T2.5, #121802, Biolegend, San Diego, CA) for 10 minutes and treated with $Pam_3CSK_4$ (Invivogen, San Diego, CA), a TLR2-specific agonist, for 10 minutes.

All values are reported as mean ± SEM.

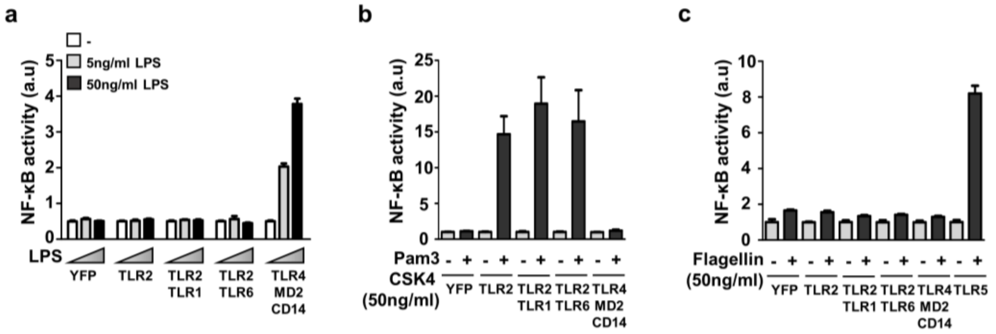


**Supplementary Figure 6. Test of reconstituted TLR system in HEK293 cells by specific agonists.**

HEK293T cells were transfected with a luciferase vector that expresses luciferase under the promoter containing NF-κB-binding elements, and various TLRs for 1 day. Then, cells were treated with LPS with indicated concentration (**a**), 50ng/ml $Pam_3CSK_4$ (**b**), or 50ng/ml recombinant Flagellin (**c**). After 24 hours later, the activity of luciferase was measured by ONE-Glo™ Luciferase Assay System according to the manufacturer's instructions. See the detail in Method.

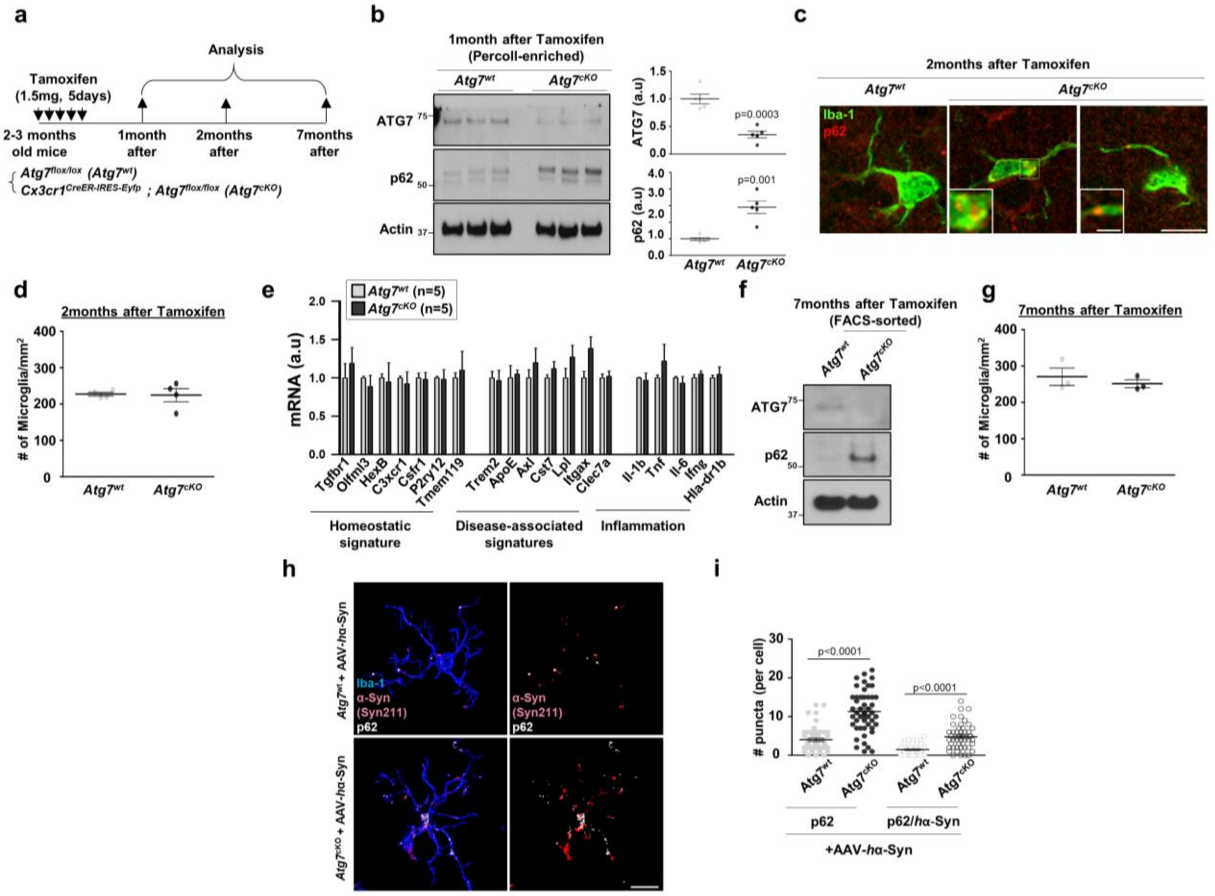


**Supplementary Figure 7. Establishment of microglia-specific *Atg7*-deficient mice.**

**(a)** Experimental plan for obtaining microglia-specific *Atg7*-deficient mice by injecting tamoxifen into *Atg7*$^{\text{flox/flox}}$ mice and *Cx3cr1*$^{\text{CreER-IRES-Eyfp}}$; *Atg7*$^{\text{flox/flox}}$ mice

**(b)** At 1-month after tamoxifen, the microglia-enriched population was collected using Percoll-gradient method and lysed for W.B using antibodies against ATG7 and p62. Actin was used as a loading control. Intensities of protein bands were quantified in the right panel. n=5 per group. *p* values were calculated by Unpaired two-tailed Student's t-test.

**(c, d)** At 2-months after tamoxifen, brain sections were fixed and stained with antibodies using Iba-1 and p62 (**c**). The number of Iba-1-positive microglia at the cortex was counted using ImageJ software (NIH, Bethesda, MD) (**d**). Scale bar, 10µm; 2µm in magnified images.

**(e)** At 2-months after tamoxifen, $CD45^{intermediate}$ and $CD11b^{high}$ microglia were isolated from brains of *Atg7*$^{flox/flox}$ mice (n=5) and *Cx3cr1*$^{CreER\text{-}IRES\text{-}Eyfp}$; *Atg7*$^{flox/flox}$ mice (n=5), and assayed for RT-qPCR using primers for genes associated with homeostatic signatures, disease-associated signatures, and inflammation.

**(f)** At 7-months after tamoxifen, pooled $CD45^{intermediate}$ and $CD11b^{high}$ microglia were isolated by flow cytometry from brains of *Atg7*$^{flox/flox}$ mice (n=2) and *Cx3cr1*$^{CreER\text{-}IRES\text{-}Eyfp}$; *Atg7*$^{flox/flox}$ mice (n=2), and assayed for W.B using antibodies against ATG7 and p62. Actin was used as a loading control.

**(g)** At 7-months after tamoxifen, Iba-1-positive microglia in the cortex were counted in $Atg7^{flox/flox}$ mice (n=3) and *Cx3cr1*$^{CreER\text{-}IRES\text{-}Eyfp}$; *Atg7*$^{flox/flox}$ mice (n=3).

**(h, i)** After 6 weeks after AAV, brain slices were fixed and stained with antibodies against human α-synuclein, p62, and Iba-1. Representative 3D reconstruction pictures of microglia containing p62 and human α-synuclein in the striatum were produced using Imaris (Bitplane, **h**). The number of p62 puncta or p62-positive/*h*α-Syn-positive puncta was quantified in microglia at striatum of *Atg7*$^{flox/flox}$ mice (n=3) and *Cx3cr1*$^{CreER\text{-}IRES\text{-}Eyfp}$; *Atg7*$^{flox/flox}$ mice (n=3) (**i**). *p* values were calculated by Mann–Whitney U test. Scale bar, 10µm.
